# Inter-token silence period spiking activity enhances selectivity of distinct groups of auditory cortical neurons to periodic and aperiodic sound sequences

**DOI:** 10.64898/2026.02.03.703433

**Authors:** A.S. Micheal, S. Bandyopadhyay

**Affiliations:** Information Processing Laboratory, Dept of E&ECE, IIT Kharagpur; Advanced Technology Development Centre, IIT Kharagpur

## Abstract

Auditory neurons, in the midbrain and beyond, detect changes in repeating acoustic patterns. Most studies focus on mechanisms underlying such sensitivity and adaptation to regularity. However, regular sound patterns are crucial in social communication, stream-segregation and grouping in different species including humans. Thus, we address cortical selectivity to periodic or aperiodic sound sequences with multiple stimulus attributes. With single unit electrophysiology and two-photon calcium imaging in anesthetized and awake mouse auditory cortex, we observe subpopulations of neurons selective to periodicity or aperiodicity that lack generalization across period-length, frequency-content or inter-token-interval durations. Comparing results with or without inter-token-interval spiking activity, shows its profound role underlying selectivity to periodic or aperiodic sequences. The whole population average rate for periodic and aperiodic stimuli is identical but not following each period and stimulus-off-period. Hence, inter-token-interval activity, post each period increases during the sequence providing information on selectivity and a prediction like signal.

**Highlights:**

- Contrary to common view, subpopulations of neurons in the auditory cortex are selective to repetitive sound patterns or periodic sound sequences and another to aperiodic ones.
- Neither population of neurons above generalizes their selectivity across properties of tokens of the sequences.
- Neural activity during the inter-token interval plays an important role in enhancing the observed selectivity.
- Post period or pre-subsequent period activity builds up during the stimulus providing a prediction like signal for periodic sequences with respect to aperiodic sequences.

## Introduction

The auditory system’s remarkable ability to detect regularity and its violations is crucial to sound perception and learning (Näätänen & Picton, 1987, Winkler et al., 2009, Okamoto et al., 2010). The midbrain to auditory cortex hierarchy needs to discriminate between complex acoustic spectral and temporal features, regular or irregular in nature, to parse intricate acoustic environments (Bregman, 1990) and alert organisms when expected patterns are disrupted (Malmierca et al., 2014; Yaron A. et al., 2012). Such responses have been explored through measures like mismatch negativity (MMN) and adaptation dynamics, revealing a general principle: neurons in the auditory pathway and beyond are tuned to detect and respond to unexpected stimuli against a backdrop of predictable sound sequences (Näätänen et al., 2007; Mehra et al., 2022a, Srivastava & Bandyopadhyay, 2020; Malmierca et al., 2009; Antunes et al., 2010; Ulanovsky et al., 2003). The idea of deviance detection have been extended to sequences of sound tokens (Mehra et al., 2022b) and have been implicated in developmental plasticity (Mehra et al., 2022a). Similarly predictable regular sequences are equally important in parsing auditory environments and play a role in social communication, experience dependent plasticity in mice and auditory stream segregation (Perrodin et al., 2023; Agarwalla et al., 2020; Agarwalla et al., 2023; Agarwalla et al., 2025; Andreou et al., 2011; Carl & Gutschalk, 2012). Similar principles of stimulus specific adaptation and deviance detection are present in vision and somatosensation, where predictable input streams heighten sensitivity to unexpected changes (Maravall et al., 2007; Adibi et al., 2013; Tang et al., 2023) and also with cross-modal sensory streams of sound and visual stimuli and vice versa (Sharma & Bandyopadhyay, 2020). Such parallels reinforce the notion that cortical circuits are shaped by a combination of adaptation and prediction, leveraging prior experience to anticipate future sensory events (Rao et al., 1999).

In the present study, we build on the above concepts to examine how mouse auditory cortical neurons encode repeated presentations of both Periodic and Aperiodic sequences using single unit recordings and with 2-photon Ca^2+^ imaging. By introducing irregularities without altering average spectral content, we directly test the sensitivity of the auditory cortex single neurons to periodic and aperiodic temporal structures at multiple time scales. We found that for all kinds of stimuli used based on spectral content and time scale and recording methodology, at least ∼5-40% of the neurons were periodic selective (PS) or Exclusive Periodic (EP) and almost similar proportion to be aperiodic selective (AS) or Exclusive Aperiodic (EA). The PS or AS nature of neurons however did not generalize across, period length in number of sound tokens, spectral content of sequences and time scales of sound tokens of the sequences. Although we find presence of both AS and PS neurons in the ACX they were not generic detectors of periodicity or aperiodicity. On investigating the nature of adaptation over the time course of the stimuli PS and AS showed distinct adaptation dynamics which could be the underlying reason for their selectivity differences. Although not generic, surprisingly, our imaging data show that at the microscale PS neurons spatially cluster together, and similarly AS neurons also spatially cluster together to within 50 μm, suggesting an underlying organization based on responses to sequences. Finally, we find that the neuronal activity in the intervening period of the tokens in the stimuli play crucial role in determining selectivity of neurons, suggesting off period effects like release from inhibition (Yarden et al., 2022), differential adaptation of inhibitory neuron types (Natan et al., 2015; Natan et al., 2017) along with synaptic adaptation leads to the two kinds of selectivity. It must be noted that, while most attention has been given to violations of regularity (Khouri & Nelken, 2015; Sharma & Bandyopadhyay, 2020; Taaseh et al., 2011) and how neurons signal such violations, as we show here selectivity to regularities is equally present in neurons of the ACX at different time scales.

## Results

### Significant proportion of ACx neurons are selective to either Periodic or Aperiodic Sequences

Periodic selectivity of single units in the mouse ACx (*N*=6 mice) was first studied with Period-3 and Period-4 stimuli with fixed-ITI of 70 ms (Figure 1A, Methods-Auditory Stimulus with fixed time scale - different Periodicities (fixed-ITI)). Results below are based on a total of 289 units, in which 138 units showed significant responses (see *Methods - significant units’ selection*). Peri-stimulus time histograms (PSTHs, Figure 2A) and the corresponding dot raster (Figure 2B) obtained from responses to the 16 stimuli (fixed-ITI stimulus set) show locking to the tokens, with strong responses initially followed by adaptation as expected. All significant units exhibited similar responses represented above (Figure 2A) for all four groups of fixed-ITI stimuli. Since the first period does not play a role in the Periodic or Aperiodic identity of the stimulus and to rule out substantial variation in responses across units, analyses is performed based on normalizing response rate of all tokens (and periods/segments) by response to the first period (or corresponding segment for Aperiodic stimuli).

**Figure 1.**
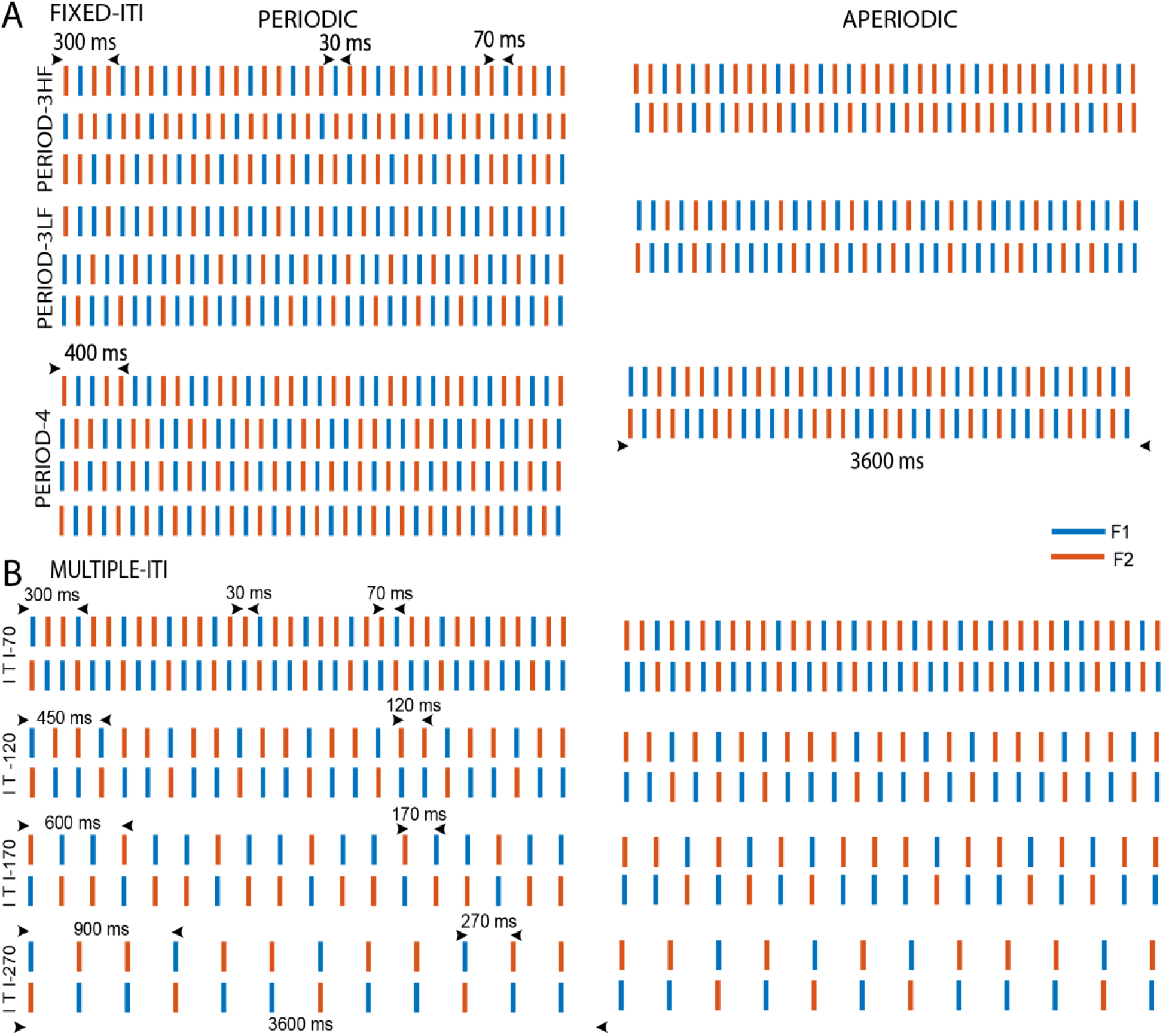
Schematic representation of the Fixed-ITI and Multiple-ITI stimulus sets. **(A)** The left panel shows periodic stimulus sets, with the first three rows representing P3HF, the next three representing P3LF, and the last four representing P4. The right panel presents the corresponding aperiodi stimuli (A3HF, A3LF, and A4) aligned with their periodic counterparts (See Methods: Auditory Stimulus with Fixed Time Scale – Different Periodicities). **(B)** The left panel displays periodic stimuli, and the right panel displays the corresponding aperiodic versions for ITIs of 70, 120, 170, and 270 ms. Blue represents *f1* and red represents *f2* in each stimulus, with the token durations and ITIs clearly marked. (See Methods: Auditory Stimulus with Multiple Timescales).

**Figure 2:**
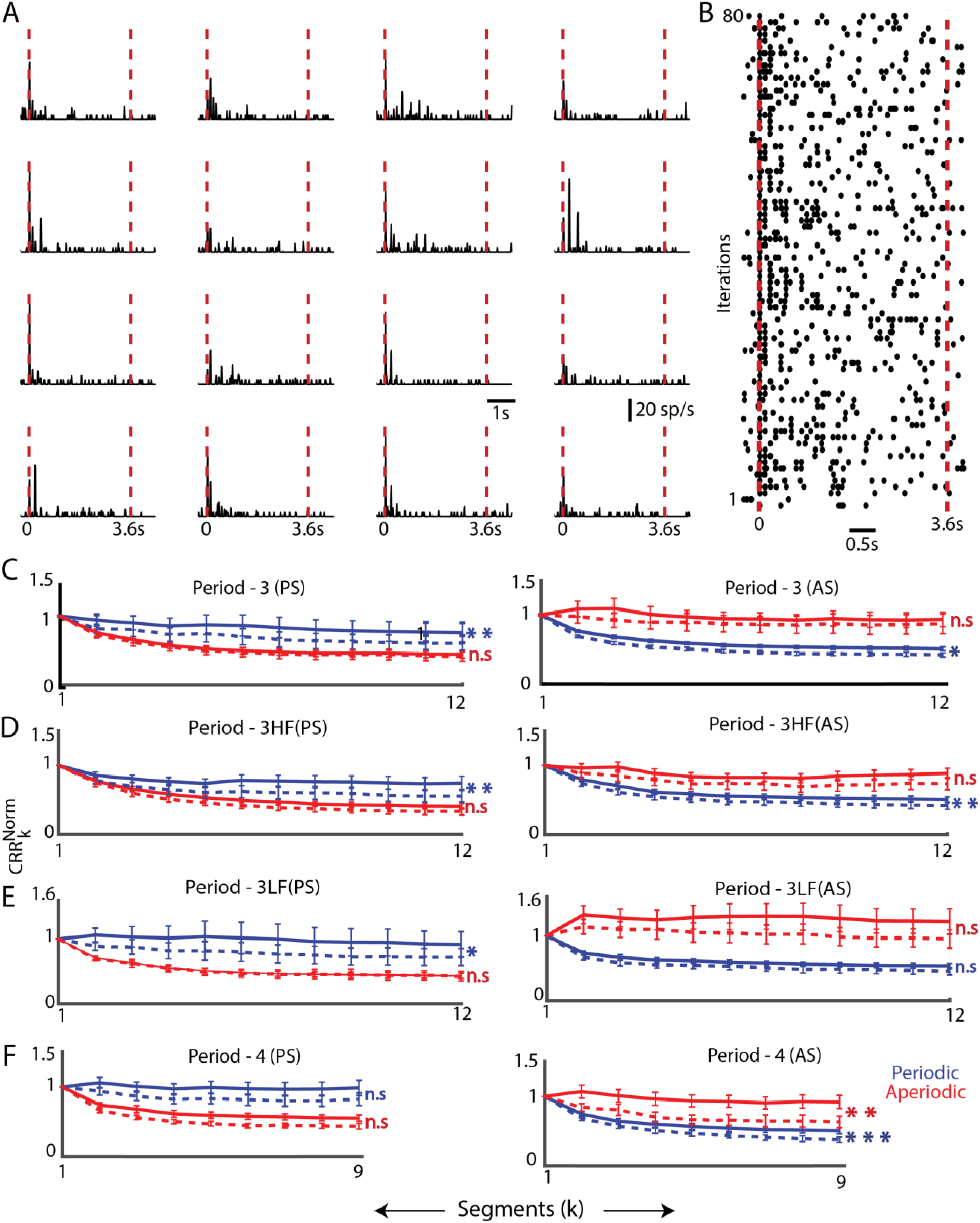
**Effect of different periodicities on Temporal Dynamics of Selective Neuronal Responses** (**A**) Example single-unit smoothed peristimulus time histogram (PSTH) using a 10 ms bin, averaged across five trials for all fixed-ITI stimuli. The red dashed vertical lines mark the onset and offset of the stimulus presentation. (**B**) Dot raster plot of the same example unit showing responses to all 16 stimuli across five trials each. The red dashed vertical lines again indicate the beginning and end of the stimulus. **(C-F) Left,** Normalized Cumulative response rates *CRR*_*k*_^*Norm*^(*withITI*) for Periodic Selective (PS) units , showing both with-ITI and without-ITI conditions. Solid blue lines represent periodic stimulus set , while solid red lines indicate aperiodic stimulus set of Period −3, Period −3HF, Period-3LF and Period-4 respectively. Dashed blue lines (without ITI) represent periodic responses and dashed red lines (without ITI) represent aperiodic responses. **Right**, Normalized Cumulative response rates *CRR*_*k*_^*Norm*^ (*withITI*) for the Aperiodic Selective (AS) units following the same format as the PS units: solid blue (periodic with ITI), solid red (aperiodic with ITI), dashed blue (periodic without ITI), and dashed red (aperiodic without ITI).

Single units however could exhibit exclusive selectivity to either the Periodic or the Aperiodic stimulus set of that group. To classify units for the above purpose, the normalized mean cumulative response rates of each stimulus (*CRR*^*Norm*^, *Methods-Normalized Cumulative response rates and Classification of single units)* for each unit was analyzed. Periodic sets (P3, P4, P3HF, P3LF) included varying numbers of Periodic stimuli (6, 4, 3, and 3 respectively), while Aperiodic sets (A3, A4, A3HF, A3LF) had 4, 2, 2, and 2 stimuli, respectively in the fixed-ITI stimulus set. Most units (>50%) were nonselective, with the proportion of PS varying between 11% and 16%, while AS units ranged from about 14% to 18% (Table 1). The single units that belong to PS will not be AS in the same stimulus group.

**Table 1.** Distribution of Periodic (PS) and Aperiodic (AS) Selectivity for Fixed-ITI Stimuli. **a-d**, Values indicate the number of single units with the corresponding percentage of single units to the total significant 138 units in parenthesis selective to Periodic and aperiodic stimulus set for the four fixed-ITI stimuli. Each column represents specific stimulus set.

| SELECTIVITY | Period-3<sup>a</sup> | Period-3LF<sup>b</sup> | Period-3HF<sup>c</sup> | Period-4<sup>d</sup> |
| --- | --- | --- | --- | --- |
| PERIODIC SELECTIVE | 16(11.7%) | 23(16.6%) | 18(13%) | 21(15%) |
| APERIODIC SELECTIVE | 24(17.7%) | 20(14.5%) | 19(14%) | 23(16.7%) |

The percentage of units which are PS and AS in each category remains same, indicating no preference to any Periodicity or frequency content. Thus, we find that units in the ACx can exhibit selectivity to Periodic sequences (with Periodicities as large as 300 and 400 ms) either with significantly higher (PS) relative responses over Aperiodic counterparts or similarly lower (AS) responses compared to the same.

### Distinct adaptation patterns for Periodic and Aperiodic stimuli

Adaptation in responses to the sequence of tokens was observed in single-unit populations and in selective subpopulations of Periodic-selective (PS) and Aperiodic-selective (AS) units. PS and AS subpopulations displayed distinct adaptation patterns across all the fixed-ITI stimulus sets for the Periodic stimuli and the Aperiodic stimuli (blue and red respectively, solid lines, Figure 2C-F, left for PS and right for AS). Strikingly lower adaptation was evident in responses of the PS to Periodic stimuli compared to responses to the Aperiodic stimuli. Similarly rate responses of the AS to the Aperiodic stimuli adapted less compared to responses to the Periodic stimuli. The above differences were quantitatively observed with exponential decay fits to the population rate response profiles over the stimuli sequences as summarized below.

The normalized cumulative response rates, *CRR*_*k*_^*Norm*^(*with − ITI*) obtained by averaging across stimuli within Periodic (P3, P4, P3HF, P3LF) and Aperiodic (A3, A4, A3HF, A3LF) sets were fitted with the exponential model (with *fit* (non-linear least squares), MATLAB). All decay time constants (τ (s)) obtained after fitting had *R*^2^ > 0.95 unless indicated otherwise. The PS units’ response to Periodic stimulus set had larger time constants (P3, τ = 0.2; P3HF, τ = 0.35; P3LF, τ = 25.6, *R*^2^ = 0.77; P4, τ = 50, *R*^2^ = 0.45; Figure 2C-F, left, blue) compared to their responses to the corresponding Aperiodic stimulus set (A3, τ = 0.5; A3HF, τ = 0.75; A3LF, τ = 0.52; A4, τ = 0.55; Figure 2C-F, left, red). Similarly, as mentioned, the AS units’ response to Periodic stimuli showed smaller time constants (P3, τ = 0.56 ; P3HF, τ = 0.62 ; P3LF, τ = 0.43; P4, τ = 0.61 ; Figure 2C-F, right side, blue) compared to their corresponding Aperiodic stimuli response time constants for all categories (A3, τ = 2.8, *R*^2^ = 0.64; A3HF, τ = 0.68, *R*^2^ = 0.74; A3LF, τ = 55, *R*^2^ = 0.12; A4, τ = 065; Figure 2C-F, right side, red). Thus, we find that the degree of adaptation in the subpopulations of neurons in response to the stimuli is related to their specific selectivity.

The Periodic or Aperiodic nature of the stimulus is inclusive of the ITI and hence activity during that period can have a role in forming selectivity to the whole stimulus set. To investigate whether the activity in the ITI period has a role we computed *CRR*_*k*_^*Norm*^(*with − ITI*) (*Methods – With-ITI and Without-ITI*). In general, we observe a lowering of the sustained nature of the response or increased adaptation (Figure 2C-F, dashed lines). The effect of the activity during the ITIs is quantified by the parameter *d* (*Methods – With-ITI and Without-ITI*) with a comparatively smaller *d* implying larger contribution of ITI period activity. The above effect was larger in case of PS in response to Periodic stimuli (P3, τ = 0.8; P3HF, τ = 0.74; P3LF, τ = 0.78 ; P4, τ = 0.83; Figure 2C-F, left, blue) compared to their response to Aperiodic stimuli (A3, τ = 0.93; A3HF, τ = 0.81; A3LF, τ = 0.98; A4, τ = 0.78; Figure 2C-F, left, red) and similarly for AS in response to Aperiodic stimuli (A3, τ = 0.93; A3HF, τ = 0.86; A3LF, τ = 0.75; A4, τ = 0.69; Figure 2C-F, right, red) compared to their responses to Periodic stimuli (P3, τ = 0.83; P3HF, τ = 0.85; P3LF,τ = 0.86 ; P4, τ = 0.75; Figure 2C-F, right, blue). Thus, as a whole in the population of units, the preference to Periodic or Aperiodic stimuli is dependent on the ongoing activity during the entire stimulus sequence – both during and between the tokens. The PS units’ response to Periodic stimulus set shows smaller time constants (P3 and A3 all) when activity during the ITIs is removed in computing the rate response profile across the cumulative segments compared to that when including the activity during the ITIs (P3, τ = 0.1 ; P3HF, τ = 0.49 ; P3LF , τ = 1.17 ; P4 , τ = 0.71; Figure 2C-F, left side, blue dashed lines).

The above units, without ITI still show stronger response to the Periodic stimulus, while their corresponding Aperiodic stimulus set response time constants for all categories do not change significantly (A3, τ = 0.55; A3HF, τ = 0.63; A3LF, τ = 0.49; A4, τ = 0.48; Figure 2C-F, left side, red dashed lines). The same is observed in the case of AS units’ responses (P3 and A3 all). The AS unit’s response to Periodic stimulus sets shows smaller time constant values (P3, τ = 0.47 ; P3HF, τ = 0.55 ; P3LF, τ = 0.35; P4, τ = 0.65 ; Figure 2C-F, right side, blue dashed lines) compared to their corresponding Aperiodic stimuli response time constants for all categories (A3 , τ = 0.75; A3HF, τ = 0.63; A3LF, τ = 24.4, *R*^2^ = 0.59; A4 , τ = 0.86; Figure 2C-F, right side, red dashed lines). The lower time constants obtained without ITI obviously also led to lower cumulative normalized responses as indicated below. A paired t-test showed significant differences in *CRR*^*Norm*^values between with-ITI and without-ITI conditions for Periodic (P3, P4, P3HF, P3LF) and Aperiodic (A3, A4, A3HF, A3LF) stimuli within PS and AS subpopulations (Figure 2C-F). The PS units showed significant differences in the responses to Periodic stimuli on considering with ITI and without ITI (P3, p<0.01; P3HF, p<0.01; P3LF, p<0.05; except for P4, p=0.09 Figure 2 C-F, left side, blue) and for no significant differences in any of the Aperiodic stimuli (A3, p=0.2; A3HF, p=0.1; A4, p=0.9; Figure 2 C-F left side, red). The AS units showed significant differences in the responses to Periodic stimuli on considering with ITI and without ITI (P3, p<0.05; P3HF, p<0.01; P4, p<0.001; Figure 2 C, D, F right side, blue) except P3LF, p=0.06; Figure 2 E right side, red) and for Aperiodic stimulus set (A4, p<0.001; Figure 2F right side, red ) except for (A3, p=0.74; A3HF, p=0.06; A3LF, p=0.31; Figure 2 C,D,E right side, red).

Thus, PS and AS neuron populations exhibited differential adaptation patterns over time (with-ITI and without-ITI) across all fixed-ITI stimulus sets. PS units adapted slower to Periodic stimuli, indicated by higher time constants, but showed faster adaptation to Aperiodic stimuli, reflecting a preference for Periodic input. Conversely, AS units adapted slower to Aperiodic stimuli and faster to Periodic stimuli. This result aligns with stimulus selectivity: PS units, selective to Periodic stimuli, exhibited higher response rates to fixed-ITI Periodic stimuli compared to Aperiodic stimuli, while AS units showed the opposite pattern. This trend persisted across Periodicity-3 and Periodicity-4 stimuli, indicating that differential adaptation profiles of units give rise to selectivity to Periodicity and Aperiodicity, regardless of differences in Periodicities or frequency content.

All plots are presented with the standard error of the mean (SEM) and the significant differences in *CRR*^*Norm*^values between with-ITI and without-ITI conditions for Periodic (P3, P4, P3HF, P3LF, Blue asterisks) and Aperiodic (A3, A4, A3HF, A3LF, Red asterisks) stimuli within PS and AS subpopulations is shown (t-test, *p<0.05, **p<0.01, ***p<0.001, ****p<0.0001, and n.s. = Not significant).

### Lack of generalization to Periodicities

In continuing our investigation of selectivity to Periodic and Aperiodic stimuli, we next wanted to assess if neurons that were Periodic selective would be selective to different Periodicities. To assess generalization across Periodicities and frequency content, we identified units overlapping between two Periodicities and two frequency contents (Table 2) within fixed-ITI stimulus sets separately for PS and AS subpopulations. For PS units, 1 unit (0.7%) exhibited selectivity across two Periodicities (Period-3 and Period-4), while 3 units (2.2%) generalized their selectivity to Periodic stimuli with different frequency content (Period-3HF and Period-3LF). For AS units, 6 units (4.4%) showed selectivity for Aperiodic stimuli across two Periodicities (Period-3 and Period-4), and 3 units (2.2%) demonstrated generalization to Aperiodic stimuli with different frequency content (Period-3HF and Period-3LF). These findings indicate that although selectivity exists for Periodic and Aperiodic stimuli, the ability of units to generalize across Periodicities and frequency content is scarce.

**Table 2:** Limited generalization of Periodic and Aperiodic selectivity to periodicities. **a,b** Values indicate the number of single units, with the corresponding percentage (in parentheses) relative to the total of 138 significant units, that are common to Period-3 and Period-4 (Column a) and to Period-3HF and Period-3LF (Column b) and are selective to the periodic and aperiodic stimulus sets for the four fixed-ITI stimuli.

| GENERALIZATION | Period-3, Period- 4<sup>a</sup> | Period-3HF, Period-3LF<sup>b</sup> |
| --- | --- | --- |
| PERIODIC SELECTIVE | 1 (0.7%) | 3 (2.2%) |
| APERIODIC SELECTIVE | 3 (2.2%) | 6 (4.4%) |

### Inter token interval activity contribute significantly to coding periodic and aperiodic stimuli

Periodic-selective (PS) and Aperiodic-selective (AS) subpopulations’ *CRR*^*Norm*^scatter responses, for each unit, with-ITI (filled circles) and without-ITI (‘+’ symbols, excluding silence of 50ms), (Figure 3A-D) were observed between the Periodic and Aperiodic stimuli in all stimulus groups. The units that are common for PS and AS in each category (Table 2) did not cluster away from the groups (Figure. 3, black and cyan symbols). The *CRR*^*Norm*^ values differ significantly between Periodic and Aperiodic stimuli across all stimulus sets with-ITI for PS units (Period-3, p<0.01; Period-3HF, p<0.0001; Period-3LF,p<0.05; Period-4, p<0.0001; blue asterisks, Figure 3A-D) and AS units (Period-3 ,p<0.001; Period-3HF, p<0.0001; Period-3LF, p<0.01; Period-4, p<0.0001; red asterisks, Figure 3A-D). This indicates that strong difference in encoding the sound sequences between Periodic and Aperiodic stimuli, when the silence periods in the sequences are considered. The *CRR*^*Norm*^ for Periodic stimuli and Aperiodic stimuli in all categories without-ITI for PS units (Period-3, p<0.05; Period-3HF, p<0.01; Period-3LF, p<0.05; Period-4, p<0.0001) and AS units (Period-3, p<0.01; Period-3HF, p<0.01; period-3LF, p<0.01; Period-4, p<0.001) also shows difference. The histograms (Figure 3A-D) along the axes show the distribution of *CRR*^*Norm*^ for each unit under all fixed-ITI stimulus sets. Filled bars represent with ITI, while dashed bars represent without ITI. Some units exhibited facilitation (outside gray boxes, Figure. 3) instead of adaptation (*CRR*^*Norm*^>1) in both PS and AS subpopulations in all groups of fixed-ITI stimulus set. PS neurons showed facilitation (Period-3, n=4; Period-3HF, n=3; Period-3LF, n=4; Period-4, n=10) for the Periodic stimulus set and AS neurons showed the same for the Aperiodic stimulus set (Period-3, n=9; Period-3HF, n=7; Period-3LF, n=8; Period-4, n=8). Units that overlap between the PS and AS categories are represented using distinct colours. In the Figure for Period-3 and Period-4, units common to the AS category are marked in black, while those common to the PS category for Period − 3 and Period - 4 are marked in blue (Figure. 3A, B). The same colour scheme is applied for overlapping units in Period-3HF and Period-3LF (Figure. 3C, D), maintaining consistency across all groups of fixed-ITI. The inclusion of silence periods modifies the unit’s response to both Periodic and Aperiodic stimulus. To assess the influence of ITI on neuronal selectivity, we directly compared the difference in *CRR*^*Norm*^ between periodic and aperiodic sequences obtained with and without ITI. For each neuron, we computed *CRR*^*Norm*^ (Periodic ,with-ITI) - *CRR*^*Norm*^ (Aperiodic ,with-ITI) and *CRR*^*Norm*^(Periodic ,without-ITI) - *CRR*^*Norm*^(Aperiodic ,without-ITI) , and compared these two quantities using a t-test within PS and AS populations. This analysis revealed a significant effect of ITI on selectivity in PS units (Period-3, p =0.35; Period-3HF, p=0.0081; Period-3LF, p = 0.059; Period-4, p =0.082) and in AS units (Period-3, p =0.871; Period-3HF, p =0.0035; Period-3LF, 0.021; Period-4, p=0.065), indicating that ITI significantly modulated selectivity for Period-3HF in both PS and AS units, and for Period-3LF in AS units, while other conditions did not show a significant effect. The scatter plot, in conjunction with the histograms (Figure. 3A-D), demonstrates that silence gaps contribute significantly to the encoding process, as evidenced by the differences in firing rate distributions and the positional shifts of units in the scatter plot. The lack of generalization across Periodicities and frequency content, combined with the influence of silence gaps in encoding, indicates the presence of distinct subpopulations of single units with selectivity to specific spectral and temporal features of the stimuli. The existence of distinct PS and AS neuronal populations suggests specialized neural mechanisms for processing different types of auditory information, accounting for variations in Periodicity and spectral content.

**Figure 3.**
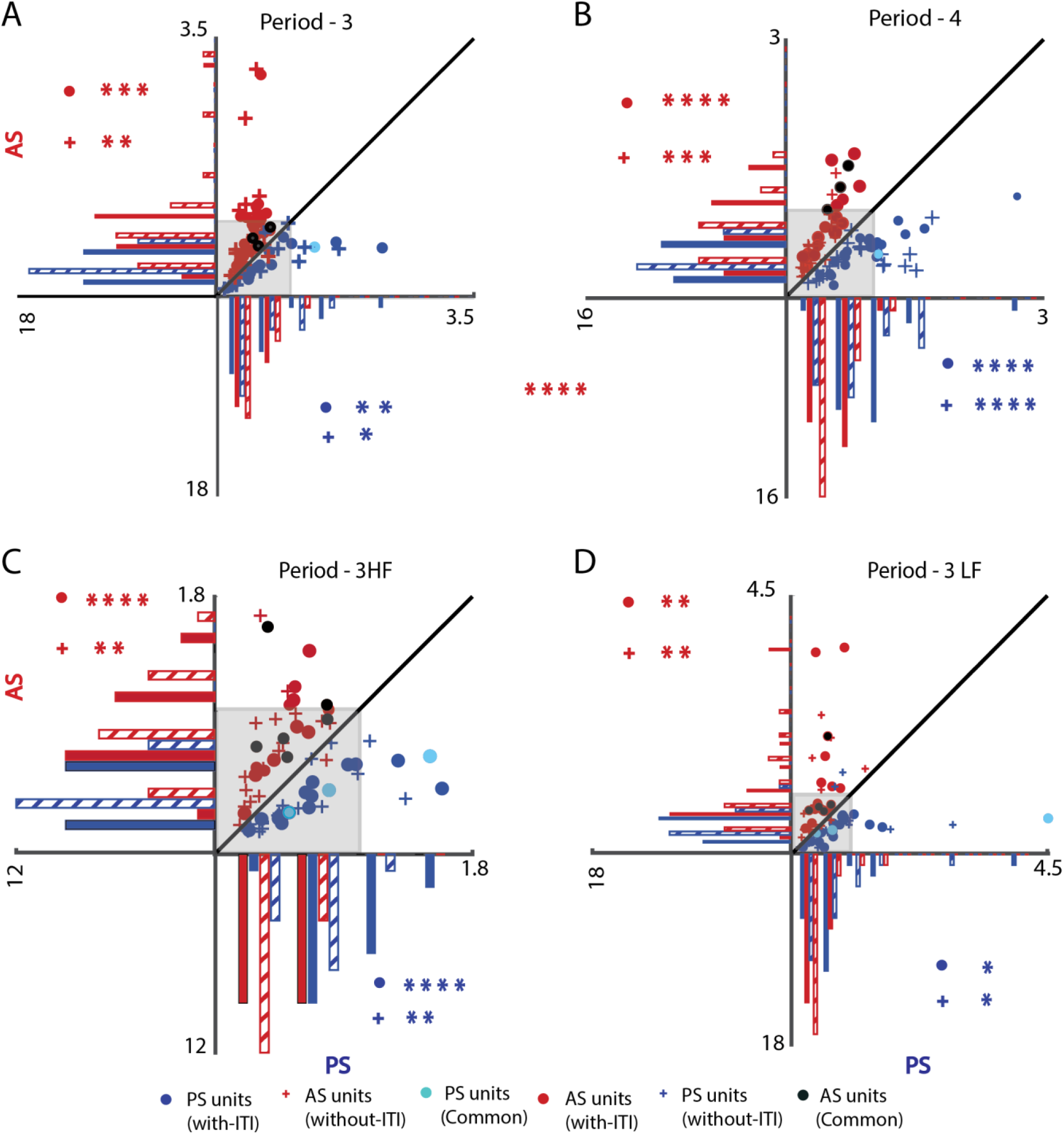
Normalized Cumulative Response Rates Highlighting Selectivity and Generalization Across Periodic and Aperiodic Stimuli. (**A-D**) Scatter plots showing the Normalized Cumulative Response Rates (*CRR*^*Norm*^) for Periodic-Selective (PS) and Aperiodic-Selective (AS) subpopulations for each unit, comparing periodic and aperiodic stimuli across all stimulus sets: Period-3, Period-4, Period-3HF, and Period-3LF. Filled circles represent responses with ITI, while ’+’ symbols represent responses without ITI. The x-axis represents the normalized cumulative firing rates for PS units, while the y-axis represents the normalized cumulative firing rates for AS units. A 45° diagonal line is shown, where units below the line are categorized as PS, and units above the line as AS. Histograms along the axes display the distribution of *CRR*^*Norm*^ with filled bars for responses with ITI and dashed bars for responses without ITI. **(A,B)** Units common between PS and AS subpopulations for Period-3 and Period-4 are marked with cyan-filled circles (PS) and black-filled circles (AS). **(C,D)** Units common between PS and AS for Period-3HF and Period-3LF are similarly marked, corresponding with the results presented in Table 2. Significant differences in *CRR*^*Norm*^is marked with asterisks: •* = with-ITI; +* = without-ITI. Colors denote subpopulations (PS, blue; AS, red). (Two-tailed t-test:, *p<0.05, **p<0.01, ***p<0.001, ****p<0.0001, and n.s. = Not significant).

### Two-photon Ca^2+^ imaging reveals distinct neuronal populations selective for Periodic and Aperiodic stimuli across different periodicities

Since single unit recordings can be biased and the single unit recordings are in anaesthetized mice, we use 2-photon Ca^2+^ imaging in awake C57BL/6J-Tg (Thy1-GCaMP6f) mice (N=3) to investigate a more unbiased population of neurons. Ca^2+^ responses to the fixed-ITI stimulus set, quantified by df/f values (Figure 4 A & B) recorded over multiple days identified 1821 significantly responding neurons out of a total of 2319 analyzed from 27 ROIs. (*Methods-Data Processing - 2-photon* 𝐶𝑎^(2+)^ *imaging - Significant cells selection).* For each stimulus set, the mean change in fluorescence (df/f) across the entire stimulus sequence and all trials was calculated for the above 1821 significantly responding neurons (Figure 4C-J, leftmost panels: blue for periodic stimuli, red for aperiodic stimuli). The mean df/f of all 1821 neurons that responded significantly were ranked from high to low based on their mean df/f values during the corresponding aperiodic stimulus (Figure 4C, E, G&I, left). Similarly, the neurons were ranked based on their df/f values during the corresponding periodic stimulus (Figure 4 D, F, H&J, left). The mean df/f of neurons with significant responses to periodic stimuli (Figure 4 C, E, G&I, middle, Table 3-Significant Periodic neurons) and that to aperiodic stimuli (Figure 4 D, F, H&J, middle, Table 3-Significant Aperiodic neurons) are retained in the process of classification of neurons. Of these remaining neurons those that also respond to Aperiodic stimuli (Figure 4C, E, G&I, right) and that to Periodic stimuli (Figure 4D, F, H&J, right) were excluded. Hence, neurons that showed significant responses only to the periodic stimulus compared to baseline were classified as Exclusive Periodic (EP), whereas those that showed significant responses only to the aperiodic stimulus compared to baseline were classified as Exclusive Aperiodic (EA). Due to the relatively low temporal resolution of calcium imaging (∼200 ms per frame) and the bidirectional nature of df/f signals, direct comparisons between periodic and aperiodic mean responses as in electrophysiology often yield differences close to chance. Therefore, neurons significantly responsive to only one stimulus type relative to baseline were classified as Exclusive Periodic (EP) or Exclusive Aperiodic (EA), whereas those responding to both were designated as Non-selective (NS). Because the proportion of “selective” neurons under the direct periodic–aperiodic comparison similar to electrophysiology is only slightly above the nominal 5% chance level, Exclusive Periodic and Exclusive Aperiodic simply indicate neurons that respond significantly to one stimulus class but not the other, rather than providing strong evidence for a robust periodic-selective population in the imaging data.

**Figure 4.**
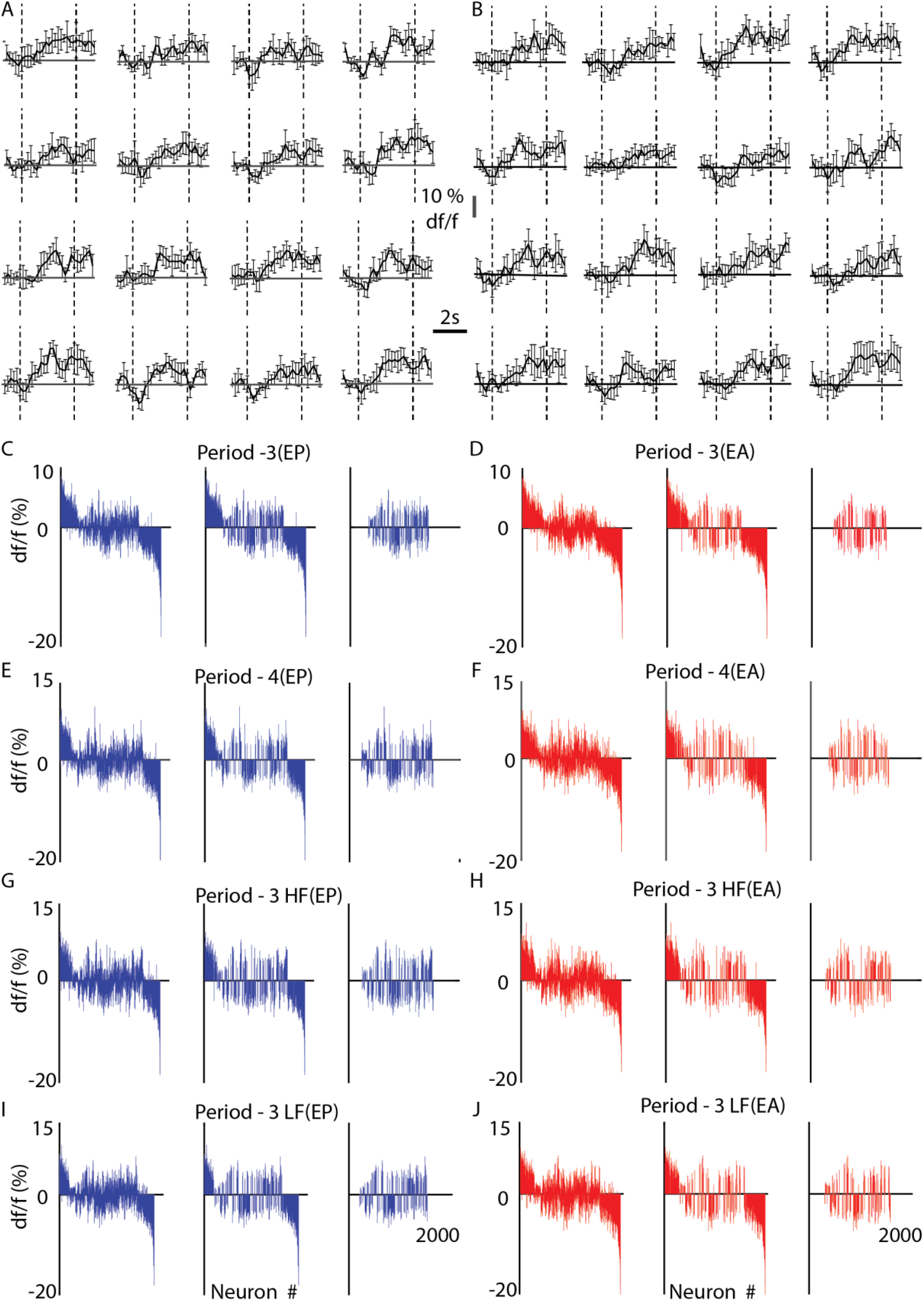
Identification of Periodic-Selective (PS) and Aperiodic-Selective (AS) neurons using 2-photon Ca²⁺ imaging in awake mice. **(A,B)** Averaged df/f traces across 5 trials in response to each of 16 fixed-ITI stimuli from two cells. The black dashed vertical lines mark the onset and offset of the stimulus presentation. **(C-J)** Mean df/f value for the entire stimulus duration across all 5 trials, shown for Periodic (blue) and Aperiodic (red) were ranked in descending order based on the neurons’ mean df/f to the opposite stimulus type. (**C-J) Left ,** Mean df/f value for all 1821 neurons identified through significant cell selection. **Middle,** Mean df/f of significant Periodic neurons to periodic stimulus set (blue) and significant Aperiodic neurons to Aperiodic stimulus set (red). **Right,** Mean df/f of Exclusive Periodic neurons to periodic stimulus set (blue) and Exclusive Aperiodic neurons to Aperiodic stimulus set (red). (see Table 3 for counts and criteria)

**Table 3.** Distribution of Significant and Selective Neuronal Units from 2-Photon Calcium Imaging in awake mice. **a,** Number of neurons that responded significantly to the different Periodic stimulus sets These neurons may or may not have also responded to Aperiodic stimuli. **b,** Number of neurons that responded significantly **only** to the Periodic stimulus sets and not to the Aperiodic stimulus sets. **c,** Number of neurons that responded significantly to the different Aperiodic stimulus sets These neurons may or may not have also responded to Aperiodic stimuli. **d,** Number of neurons that responded significantly **only** to the Aperiodic stimulus sets and not to the Periodic stimulus sets. Percentages shown in parentheses represent the proportion of neurons relative to the total of 1,821 significant neurons.

| SELECTIVITY | SIGNIFICANT PERIODIC NEURONS <sup>a</sup> | EXCLUSIVE PERIODIC (EP) <sup>b</sup> | SIGNIFICANT APERIODIC NEURONS <sup>c</sup> | EXCLUSIVE APERIODIC (EA) <sup>d</sup> |
| --- | --- | --- | --- | --- |
| Period-3 | 852(46.8%) | 304(16.7%) | 723(39.7%) | 175(9.6%) |
| Period-4 | 722(39.7%) | 354(19.4%) | 519(28.5%) | 151(8.3%) |
| Period -3HF | 629(34.5%) | 305(16.8%) | 523(28.7%) | 199(10.9%) |
| Period -3LF | 619(34%) | 296(16.3%) | 514(28.2%) | 191(10.5%) |

Compared to baseline, 35% to 47% of neurons exhibited significant responses to Periodic stimulus sets (P3, P4, P3HF, P3LF), while 28% to 40% responded significantly to Aperiodic stimulus sets (A3, A4, A3HF, A3LF) within fixed-ITI conditions (Table 3, Periodic and Aperiodic significant neurons). Neurons exhibiting responsiveness to both periodic and aperiodic stimuli within the same periodicity set were excluded from consideration as selective *(Methods-Data Processing - 2-photon* 𝐶𝑎^(2+)^ *imaging-Classification of neurons)*. Exclusive Periodic (EP) neurons accounted for 16% to 20%, and Exclusive Aperiodic (EA) neurons accounted for 8% to 11% (Table 3, Exclusive periodic and aperiodic neurons), indicating distinct neuronal subpopulations with clear preferences for Periodic or Aperiodic stimuli across varying periodicities and frequency content.

We observe that neurons with strong responses to Periodic stimuli tended to respond strongly to Aperiodic stimuli (both positive and negative magnitudes) and thus the EP and EA units all had intermediate df/f values (Figure 4C-J). Despite differences in Periodicities and frequency content, the proportions of EP and EA neurons remain relatively consistent, emphasizing the robustness of selectivity in these subpopulations. Thus, with imaging also we observe similar proportion of EP and EA neurons, as observed with single unit recordings.

As with single units, analysis of overlapping exclusive neurons revealed that 4.5% of the total population, were exclusive periodic and 1% were exclusive aperiodic across Period-3 and Period-4 (Table 4). For frequency content generalization, neuronal selectivity differed across spectral combinations, with 3% and 0.12% selective neurons observed for Period-3HF and Period-3LF (Table 4), respectively.

**Table 4.** Limited generalization across Periodicities and frequency content from 2-Photon Calcium Imaging. **a,** Number of neurons that are common to Period-3 and Period-4, with their selectivity shown in rows. **b,** Number of neurons that are common to Period-3HF and Period-3LF, with their selectivity shown in rows. Percentages shown in parentheses represent the proportion of neurons relative to the total of 1,821 significant neurons.

| GENERALIZATION | Period- 3, Period- 4 <sup>a</sup> | Period- 3HF, Period- 3LF <sup>b</sup> |
| --- | --- | --- |
| EXCLUSIVE PERIODIC (EP) | 81(4.5%) | 55(3%) |
| EXCLUSIVE APERIODIC (EA) | 18(1%) | 23(0.12%) |

The data demonstrate limited generalization across Periodicities and frequency content, highlighting the specificity of neuronal responses to stimulus types. This distinct selectivity, dependent on Periodicity and spectral features, aligns with patterns observed in single-unit electrophysiology recordings, where similar near absence of generalization and selectivity were noted. Thus, in awake conditions and relatively unbiased sampling with Ca^2+^ imaging we observe the same answers in terms of selectivity.

### Functional connectivity using noise correlations

Noise correlations, which reflect the shared variability in responses of neuron pairs, play a crucial role in encoding and understanding the functional organization of neural circuits. Among the 1821 significant neurons, those that did not belong to the EP or EA categories were classified as non-selective (NS). We computed pair-wise noise correlations (see-*methods-Noise correlations*), considering all combinations: EP-EP, EA-EA, NS-NS, EP-EA, and NS-EP. The mean pairwise noise correlations were calculated separately for all Periodic and Aperiodic stimuli as a function of distance between the pairs (Figure. 5, blue and red respectively). The 95% confidence intervals were estimated based on bootstraps (n = 500). The mean noise correlation values based on at least 10 pairs (Figure. 4, inverted histograms) at any distance was considered significant if the mean was different from 0 (t-test, p<0.05). Correlations were considered significantly different between Periodic and Aperiodic stimuli if their confidence intervals did not overlap.

**Figure 5.**
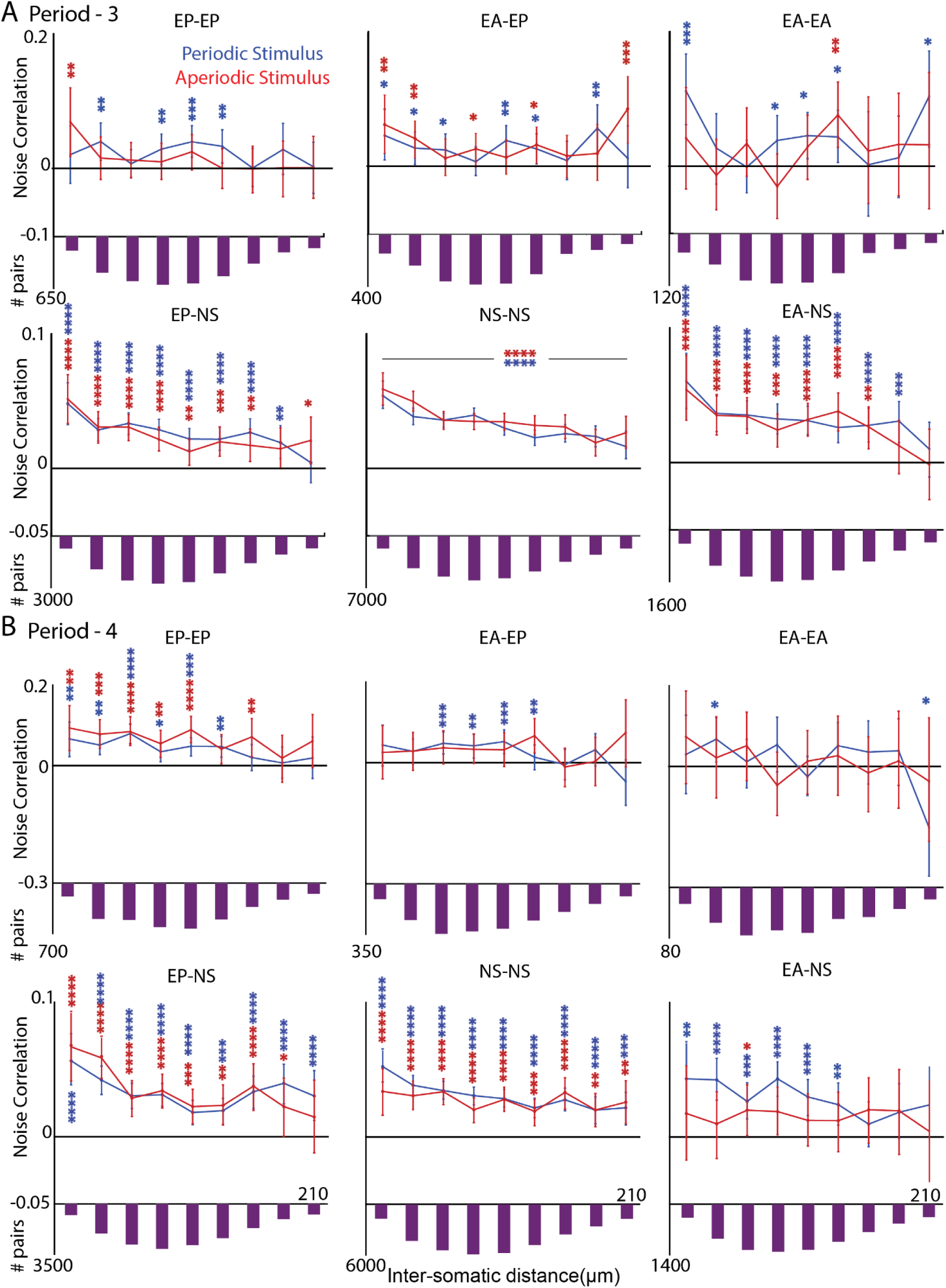
Distance-dependent noise correlations across different neuronal categories for Periodic and Aperiodic stimulus sets. Mean pairwise noise correlations as a function of inter-neuronal distance for Period-3 (**A**) and Period-4 (**B**) stimulus sets. Noise correlations were computed separately for Periodic (blue) and Aperiodic (red) stimuli, across 500 bootstrap iterations. Error bars represent the 95% confidence intervals, and individual data points indicate the mean noise correlation for each imaging field at a distance (∼23.48 μm per bin). Statistical significance was assessed using a t-test (*p* < 0.05) to test whether the mean correlation at each distance differed from zero for the periodic stimuli (blue) and aperiodic stimuli(red). (two-tailed t-test, *p<0.05, **p<0.01, ***p<0.001, ****p<0.0001, and n.s = Not significant). The inverted bar plots below each line graph indicate the number of neuron pairs in each distance bin used to compute the corresponding mean noise correlation values for each neuronal category (EP–EP, EA–EP, NS–NS, etc.).

Distinct noise correlation patterns were observed across all neuronal combinations: EP-EP, EA-EA, and EA-EP (Figure. 5A&B, top row). However, noise correlations between NS neurons and any type (EP-NS, NS-NS and EA-NS) exhibited a general decrease with increasing distance for both periodic and aperiodic stimulus sets (Figure 5 A&B; bottom row). Noise correlations in EP-EP neurons remain relatively stable across distances for Period-3 (Figure. 5A, top left), with consistently stronger functional connectivity observed for Periodic stimuli compared to Aperiodic stimuli. EA-EA neurons demonstrate moderately strong noise correlations for Periodic stimuli, particularly at larger distances (Figure. 5A, top, left). In contrast, EA-EP of Period-3 interactions display weak and inconsistent noise correlations for both Periodic and Aperiodic stimuli, indicating limited functional connectivity between these neuronal categories (Figure. 5A, top, middle). Period-4, EP-EP neurons exhibit strong and stable functional connectivity for both Periodic and Aperiodic stimuli, although connectivity weakens as the distance increases (Figure. 5B, top left). Period-4, EA-EP neurons show variable noise correlations, fluctuating around zero for Aperiodic stimuli, while some connectivity is observed at certain distances for Periodic stimuli (Figure. 5B, top middle). EA-EA neurons do not exhibit much functional connectivity, for either Periodic or Aperiodic stimuli.

### Neurons with selectivity for a stimulus type spatially cluster together

The probability of encountering a certain type (EP or EA) of neuron within a specific distance of a particular type (EP or EA) of neuron was computed across all pairs of types in each fixed-ITI stimulus set (*Methods - Spatial Arrangement*). Mean probabilities of presence of different neuron pairs (e.g., EP-EP, EP-EA, EA-EP, EA-EA) at different distances were considered significantly different if their 95% confidence intervals did not overlap.

EP neurons consistently had higher probability of being in proximity to other EP neurons across small distances (up to ∼50 μm), than by chance, indicating stable spatial clustering across all stimulus categories (Figure 6A-D, left panel, blue) and lesser probability of proximity to the EA neurons categories (Figure 6 A-D, left panel, red). Similarly, EA neurons also had higher than chance level of being near other EA neurons at short distances (50 μm, Figure 6A-D, right, red) and were near EP neurons only at chance levels. Thus, neurons of the same selectivity type had higher probability to be within 50 μm of each other, whereas those of opposite types did not spatially cluster together significantly. The above results coupled with noise correlation based connectivity results (Figure. 5) shows that there is a functional micro organization of EA and EP type neurons in the ACx. While heterogeneous in general, the organization can be thought to be like a sea of mainly NS and non-responding neurons with tight functionally coupled EA and EP neuron clusters with 100 μm diameter.

**Figure 6.**
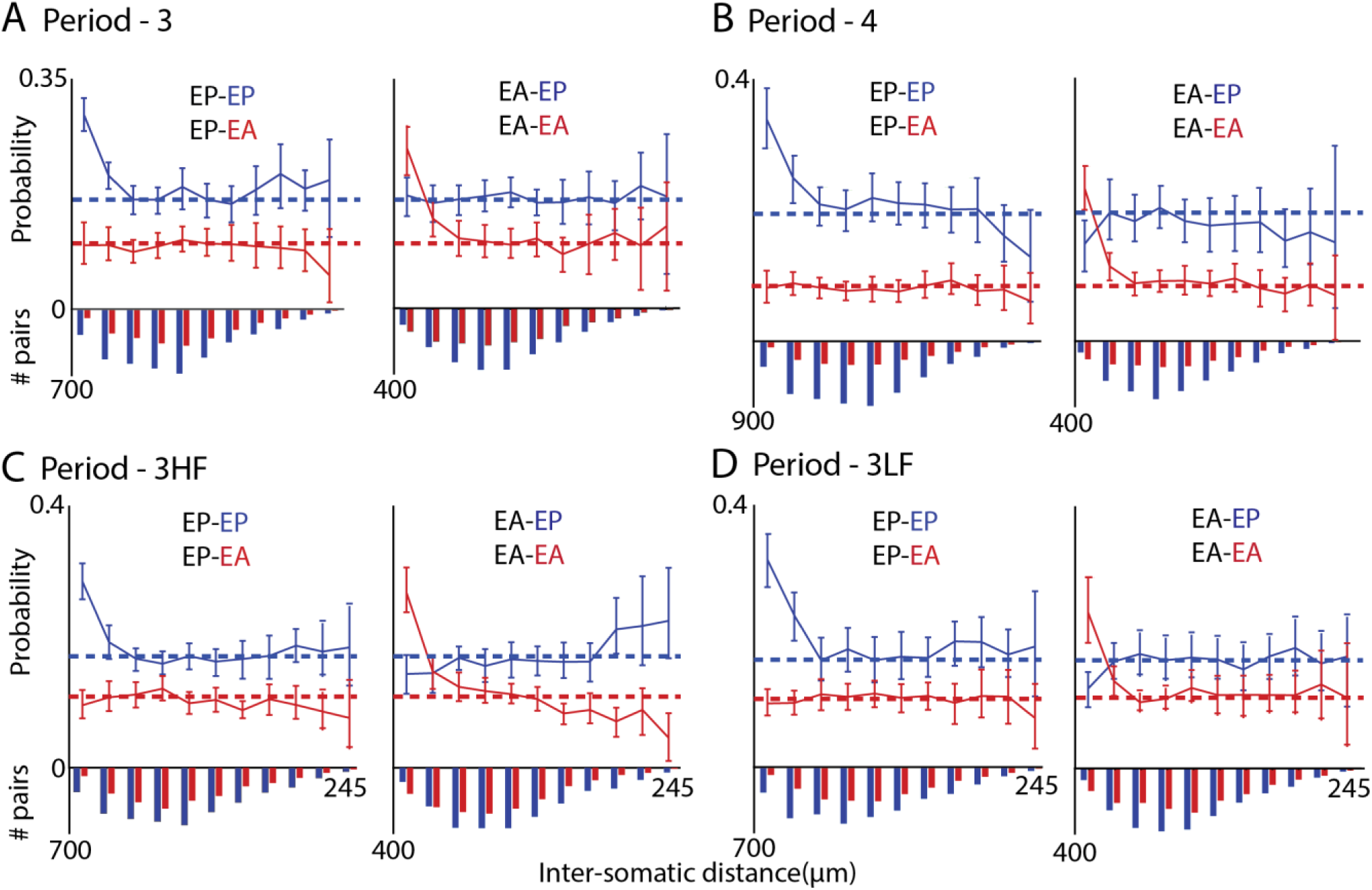
Spatial clustering of neurons based on stimulus selectivity. (**A–D) Left**, Mean probability of encountering a PS neuron (blue) or AS neuron (red) within a distance (∼23.48 μm per bin) from a reference PS neuron. **Right**, Mean probability of encountering a EP (blue) or EA (red) neuron from a reference EA neuron. Dashed horizontal lines indicate the mean probability across all distances for each pairing type.

Inverted bar plots below each panel indicate the number of neuron pairs in each distance bin: EP–EP and EA–EP in blue, EP–EA and EA–EA in red.

### Distinct Neural Populations for Periodic and Aperiodic Auditory Input: Impact of Inter-Token-Interval (ITI)

The effect of varying it is (Inter-Token Intervals) in sound sequences was analyzed using a multiple-ITI stimulus set (Figure 1B, *Methods - Auditory Stimulus with Multiple Timescales*). Responses of ACx of 575 single units (N=5 mice) to the above stimuli were collected of which 306 were deemed responsive (Methods*: Responsive Units’ Selection - Electrophysiology*). The duration of a single period varied with the ITI, with periods of 300 ms, 450 ms, 600 ms, and 900 ms corresponding to ITIs of 70 ms, 120 ms, 170 ms, and 270 ms, respectively, yielding 12, 8, 6, and 4 periods or segments respectively, per stimulus. Dot raster (Figure 7A) of all trials and the peri-stimulus time histograms (PSTHs, Figure 7B) from responses to the 16 stimuli in the multiple-ITI stimulus set demonstrated locking to token, with strong initial responses followed by adaptation. All significant units exhibited similar response patterns (Figure 7B) across all four groups of multiple-ITI stimuli.

**Figure 7.**
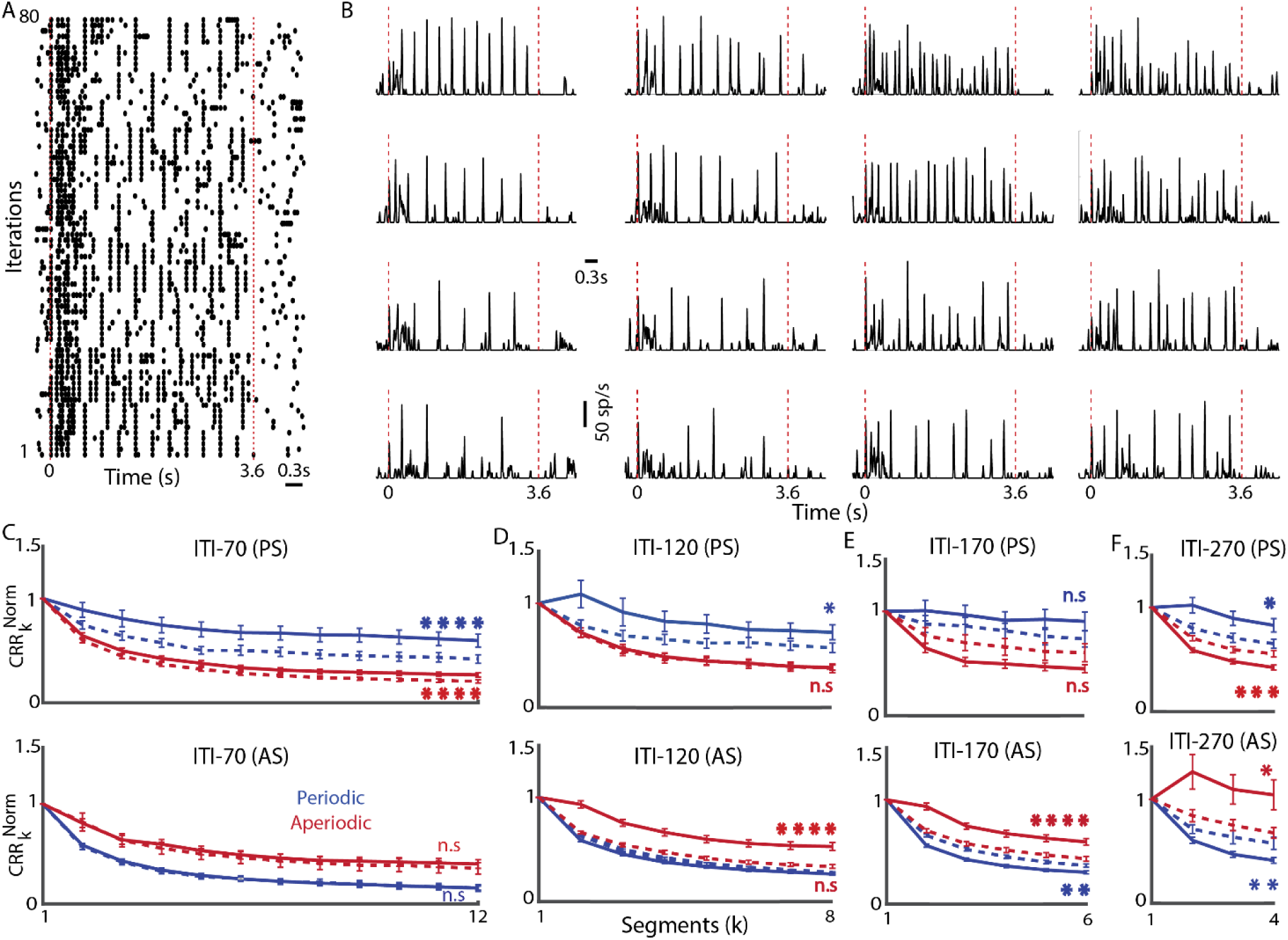
Effect of Inter-Trial Interval (ITI) on Temporal Dynamics of Selective Neuronal Responses. (**A**) Dot raster plot of the same example unit showing responses to all 16 stimuli across five trials each. The red dashed vertical lines again indicate the beginning and end of the stimulus. **(B)** Example single-unit smoothed peristimulus time histogram (PSTH) using a 10 ms bin, averaged across five trials for all multiple-ITI stimuli. The red dashed vertical lines mark the onset and offset of the stimulus presentation. **(C-F) Top,** Normalized Cumulative response rates *CRR*_*k*_^*Norm*^ (with − ITI) for Periodic Selective (PS) units, showing both with-ITI and without-ITI conditions. Solid blue lines represent periodic stimulus set, while solid red lines indicate aperiodic stimulus set of ITI-70, ITI-120, ITI-170, and ITI-270, respectively. Dashed blue lines represent periodic responses (without-ITI) and dashed red lines represent aperiodic responses (without-ITI). **Bottom,** Normalized Cumulative response rates *CRR*_*k*_^*Norm*^(with − ITI) for the Aperiodic Selective (AS) units following the same format as the PS units: solid blue (periodic with ITI), solid red (aperiodic with ITI), dashed blue (periodic without ITI), and dashed red (aperiodic without ITI). All plots are presented with the standard error of the mean (SEM) and the significant differences in *CRR*^*Norm*^ values between with-ITI and without-ITI conditions for Periodic (Blue stars) and Aperiodic (Red stars) stimuli within PS and AS subpopulations is shown (two-tailed t-test, *p<0.05, **p<0.01, ***p<0.001, ****p<0.0001, and n.s. = Not significant).

Single units were classified as Periodic or Aperiodic selective (PS and AS) based on normalized cumulative firing rates for each stimulus (*CRR*^*Norm*^) (*Methods-Normalized Cumulative response rates*) as done in the fixed time scale stimuli. Distinct groups of single units selective for Periodic and Aperiodic stimuli were observed for each ITI sequence, highlighting the segregation of neuronal responses based on stimulus timescale and pattern (Table 5). The Periodic-selective (PS) units comprised approximately 5–18% of the total 306 responsive units, while the Aperiodic-selective (AS) units accounted for around 11–38%.

**Table 5.** Distribution of Periodic (PS) and Aperiodic (AS) Selectivity of single units for Multiple-ITI Stimuli. **a-d,** Values indicate the number of single units, with the corresponding percentage (in parentheses) relative to the total of 306 significant units, that are selective to the periodic and aperiodic stimulus sets. Each column represents a specific stimulus set for the four multiple-ITI stimuli.

| SELECTIVITY | ITI-70<sup>a</sup> | ITI-20<sup>b</sup> | ITI-170<sup>c</sup> | ITI-270<sup>d</sup> |
| --- | --- | --- | --- | --- |
| Periodic Selective (PS) | 35(11.4%) | 21(6.8%) | 15(5%) | 55(18%) |
| Aperiodic Selective (AS) | 36(11.6%) | 109(35.6%) | 116(38%) | 41(13.4%) |

The observed variation in the percentage of neurons classified as Periodic-selective (PS) and Aperiodic-selective (AS) across different ITIs suggests that neuronal selectivity depends strongly on the temporal structure of the stimuli. A significant proportion of neurons, particularly at intermediate ITIs (120 and 170 ms), are classified as Aperiodic-selective. Whereas, the PS units were more in the shortest and longest it is. The above suggests a differential pattern of timescale preference in the population of PS and AS units.

### Lack of generalization to various ITI

Earlier we observed that PS and AS units inherently did not generalize to other period lengths or frequency content of the stimuli. Here we seek to see if the observed neuronal selectivity to periodicity or aperiodicity generalizes across varying inter-token intervals (ITIs) or timescales of the stimuli. We obtained the overlap of the PS and AS units across the different multiple-ITI stimulus sets. The overlapping units across all ITIs’ (Table 6) indicates a lack of generalization across ITIs, with no units classified as AS overlapping and PS overlapping across all ITIs. The highest percentage of units showing generalization for any pair or combination occurs in the AS population between ITI-120 and ITI-170(15%). In contrast, the overlap percentages for Periodic-selective units are consistently low, reflecting limited generalization.

**Table 6.** Lack of generalization across Multiple-ITI stimulus sets. **a–f.** Values indicate the number of single units, with the corresponding percentage (in parentheses) relative to the total of 306 significant units, that are common across the specified combinations of multiple-ITI stimuli and are selective to either the periodic or aperiodic stimulus sets (as indicated in rows). **g.** Values indicate the number of single units that are common across all ITI conditions and are selective to either the periodic (PS) or aperiodic (AS) category.

| GENERALIZATION | ITI-70<br>ITI-120<sup>a</sup> | ITI-70<br>ITI-170<sup>b</sup> | ITI-70<br>ITI-270<sup>c</sup> | ITI-120<br>ITI-170<sup>d</sup> | ITI-120<br>ITI-270<sup>e</sup> | ITI-170<br>ITI-270<sup>f</sup> | All ITI<sup>g</sup> |
| --- | --- | --- | --- | --- | --- | --- | --- |
| Periodic Selective<br>(PS) | 3<br>(1%) | 2<br>(0.65%) | 3<br>(1%) | 2<br>(0.7%) | 4<br>(1.3%) | 8<br>(2.6%) | 0 |
| Aperiodic Selective<br>(AS) | 10<br>(3.3%) | 11<br>(3.6%) | 4<br>(1.3%) | 46<br>(15%) | 15<br>(4.9%) | 12<br>(4%) | 0 |

### Distinct adaptation patterns for different ITI

Adaptation was observed in the subpopulations of neurons selective for Periodic and Aperiodic stimuli across all ITIs. Like the fixed-ITI stimulus set, the PS and AS subpopulations displayed distinct adaptation patterns across all the multiple-ITI stimulus sets for the Periodic stimuli and the Aperiodic stimuli (blue and red respectively, solid lines, Figure 7C-F, left for PS and right for AS).

The PS units showed lower adaptation to Periodic stimuli compared to responses to the Aperiodic stimuli, while the AS adapted less to the Aperiodic stimuli compared to responses to the Periodic stimuli. These differences were quantitatively analysed using exponential decay fits to the population rate response profiles across the stimulus sequences, as summarized below.

The normalized cumulative response rates, *CRR*_*k*_^*Norm*^(𝑤𝑖𝑡ℎ − 𝐼𝑇𝐼) obtained by averaging across stimuli within Periodic (ITI-70P, ITI-120P, ITI-170P, ITI-270P) and Aperiodic (ITI-70A, ITI-120A, ITI-170A, ITI-270A) sets each set comprising of LF and HF (one stimulus each) were fitted exponential model (with *fit* (non-linear least squares), MATLAB). All decay time constants (τ (s)) obtained after fitting had *R*^2^ > 0.95 unless indicated otherwise.

As with the fixed ITI stimulus set the PS unit’s response to Periodic stimuli showed larger time constants (ITI-70P, τ = 0.90; ITI-120P, τ = 3.2, *R*^2^ = 0.87; ITI-170P, τ = 21, *R*^2^ = 0.86; ITI-270P, τ = 13.4, *R*^2^ = 0.82 ; Figure 7C-F, top, blue) compared to their responses to the corresponding Aperiodic stimulus set (ITI-70A, τ = 0.51 ; ITI-120A, τ = 0.75; ITI-170A, τ = 0.58 ; ITI-270, τ = 0.76 ; Figure. 7C-F, top, red). Similarly, the AS unit’s response to Periodic stimuli also shows larger time constants (ITI-70P, τ = 0.51; ITI-120P, τ = 0.6; ITI-170P, τ = 0.62 ; ITI-270P, = 0.88 ; Figure 7C-F, bottom, blue) compared to their corresponding Aperiodic stimuli response time constants (ITI-70A, τ = 0.76; ITI-120A, τ = 1.53; ITI-170A , τ = 2.3 ; ITI-270A, τ = 202, *R*^2^ = 0.003 ; Figure. 7C-F, bottom, red). Thus, the degree of adaptation in neuronal subpopulations was found to depend on their specific selectivity independent of ITI. Neurons within the same preference group exhibited relatively sustained responses to their preferred stimulus type.

To investigate whether the activity in the ITI period has a role we computed *CRR*_*k*_^*Norm*^(*with − ITI*)(*Methods – with-ITI and without-ITI*). In general, we observe a lowering of the sustained nature of the response or increased adaptation for the Periodic stimulus set for PS (Figure 7C-F, blue dashed lines, top) and Aperiodic stimulus set for AS (Figure 7C-F, red dashed lines, bottom). However, as the ITI increases from 70 ms to 270 ms, we observed a strikingly opposite phenomenon of reduced adaptation to the less selective stimulus set, Aperiodic stimulus set responses in PS (Figure 7C-F, red dashed lines, top) and Periodic stimulus set responses in AS (Figure 7C-F, blue dashed lines, bottom). The PS unit’s response to Periodic stimuli shows larger time constants without-ITI (ITI-70P, τ = 0.6 ; ITI-120P, τ = 0.66; ITI-170P, τ = 2; ITI-270P, τ = 1.32 ; Figure 7C-F, blue dashed lines, top) compared to their Aperiodic stimuli response time constants for all categories (ITI-70A, τ = 0.49 ; ITI-120A, τ = 0.68; ITI-170A, τ = 0.78; ITI-270A, τ = 0.9; Figure 7C-F, red dashed lines, top). The AS units response to Periodic stimuli shows smaller time constants (ITI-70P, τ = 0.48 ; ITI-120P,τ = 0.73; ITI-170P, τ = 0.84; ITI-270P, τ = 0.87; Figure 7C-F, blue dashed lines, bottom) compared to their corresponding Aperiodic stimuli response time constants (ITI-70A, τ = 0.78 ; ITI-120A, τ = 0.74; ITI-170A, τ = 0.89; ITI-270A, τ = 2.15 ; Figure 7C-F, red dashed lines, top).

The effect of the activity during the ITIs quantified by parameter, *d* (M*ethods- with-ITI and without-ITI*) was more in case of Periodic stimuli responses of PS (ITI-70P, τ = 0.70; ITI-120P, τ = 0.79; ITI-170P, τ = 0.81 ; ITI-270P, τ = 0.78; Figure 7C-F, top, blue) compared to their response to Aperiodic stimuli (ITI-70A, τ = 0.77; ITI-120A, τ = 0.98; ITI-170A, τ = 1.34; ITI-270A, τ = 1.3; Figure 7C-F, top, red) and Aperiodic stimuli responses of AS (ITI-70A, τ = 0.91; ITI-120A ,τ = 0.64; ITI-170A, τ = 0.73; ITI-270A, τ = 0.65; Figure 7C-F, bottom, red) compared to their responses to Periodic stimuli (ITI-70P, τ = 1; ITI-120P, τ = 1.03; ITI-170P, τ = 1.21 ; ITI-270P, τ = 1.4; Figure 7C-F, bottom, blue). The PS units showed significant differences in the responses to Periodic stimuli between *CRR*^*Norm*^(with-ITI) and *CRR*^*Norm*^(without-ITI), (ITI-70P, p<0.0001; ITI-120P, p=0.03; ITI-270P, p=0.02; Figure 7C,D,F, top, blue), and Aperiodic stimuli (ITI-70A, p<0.0001 ITI-270A, p<0.001; Figure 7C,E,F, top, red), while there was no significant differences in ITI-170P, 𝑝 = 0.17 (Figure 7E, top, blue) ;ITI-120A, p=0.86 and ITI-170A, p=0.1; (Fig. 7D,E top, red). Similarly AS units showed significant differences to Aperiodic stimuli (ITI-120A, p<0.0001; ITI-170A, p<0.0001; ITI-270A, p=0.018; Figure 7D,E,F, bottom, red) and Periodic stimuli (ITI-170P, p<0.01; ITI-270A, p<0.01; Figure 7E,F, bottom, blue) and no significant differences in responses to with-ITI and without-ITI (ITI-70A, p =0.10 ; Figure 7C bottom, red) and (ITI-70P, p=0.99; ITI-120P, p=0.21 ; Figure 7C,D, bottom, blue).

Overall, in the PS and AS subpopulation of units, the preference for periodic or aperiodic stimuli depended on the ongoing activity throughout the stimulus sequence, both during and between the tokens. The effect of activity in the ITI interval increased with the ITI.

### Neuronal activity during longer ITI enhances Periodic and Aperiodic discrimination

As with the fixed-ITI stimulus set (Figure. 3), we observed the scatter of normalized cumulative response rates *CRR*^*Norm*^ with ITI (filled circles) and without ITI (‘+’ symbols, excluding silence duration corresponding to each ITI), for Periodic-selective (PS, blue) and Aperiodic-selective (AS, red) subpopulations (Figure 8A-D). Statistical comparisons reveal significant differences in *CRR*^*Norm*^between Periodic and Aperiodic stimuli in all ITI categories when considering the whole sequence (with-ITI). For PS units (ITI-70, p<10^-6^; ITI-120, p<10^-4^; ITI-170, p<10^-4^; ITI-270, p<10^-6^) and AS units (ITI-70, p<10^-6^; ITI-120, p<10^-6^; ITI-170, p<10^-6^; ITI-270, p<10^-4^). These results show marked difference in encoding sound sequences between Periodic and Aperiodic stimuli when the entire sequence is considered. When excluding ITI activity (omitting the corresponding silence periods), the differences persist but show some variation. For PS units, significant differences remain for (ITI-70, p<10^-5^; ITI-120, p=0.03; ITI-270, p=0.02) except for ITI-170, p=0.26 and AS units (ITI-70, p=0.001; ITI-120, p<10^-5^; ITI-170, p=0.02) also show differences except for ITI-270, p=0.19. This suggests that the exclusion of silence intervals reduces the distinction between Periodic and Aperiodic encoding, particularly for longer ITIs. The histograms (Figure 8A-D) along the axes show the distribution of *CRR*^*Norm*^ for each unit under all multiple-ITI stimulus sets. Filled bars represent with-ITI, while dashed bars represent without-ITI cases. Although most single units in each category exhibited differential adaptation based on the category and stimulus type, some neurons displayed facilitation (*CRR*^*Norm*^>1, outside gray boxes, Figure. 8). The number of units exhibiting facilitation in response to Periodic stimuli among PS units increased with ITI (ITI-70, n=3; ITI-120, n=3; ITI-170, n=7; ITI-270, n=11) and similarly that for Aperiodic stimuli among AS units (ITI-70, n=1; ITI-120, n=8; ITI-170; n=15; ITI-270, n=13; Figure 8 A-D).

**Figure 8.**
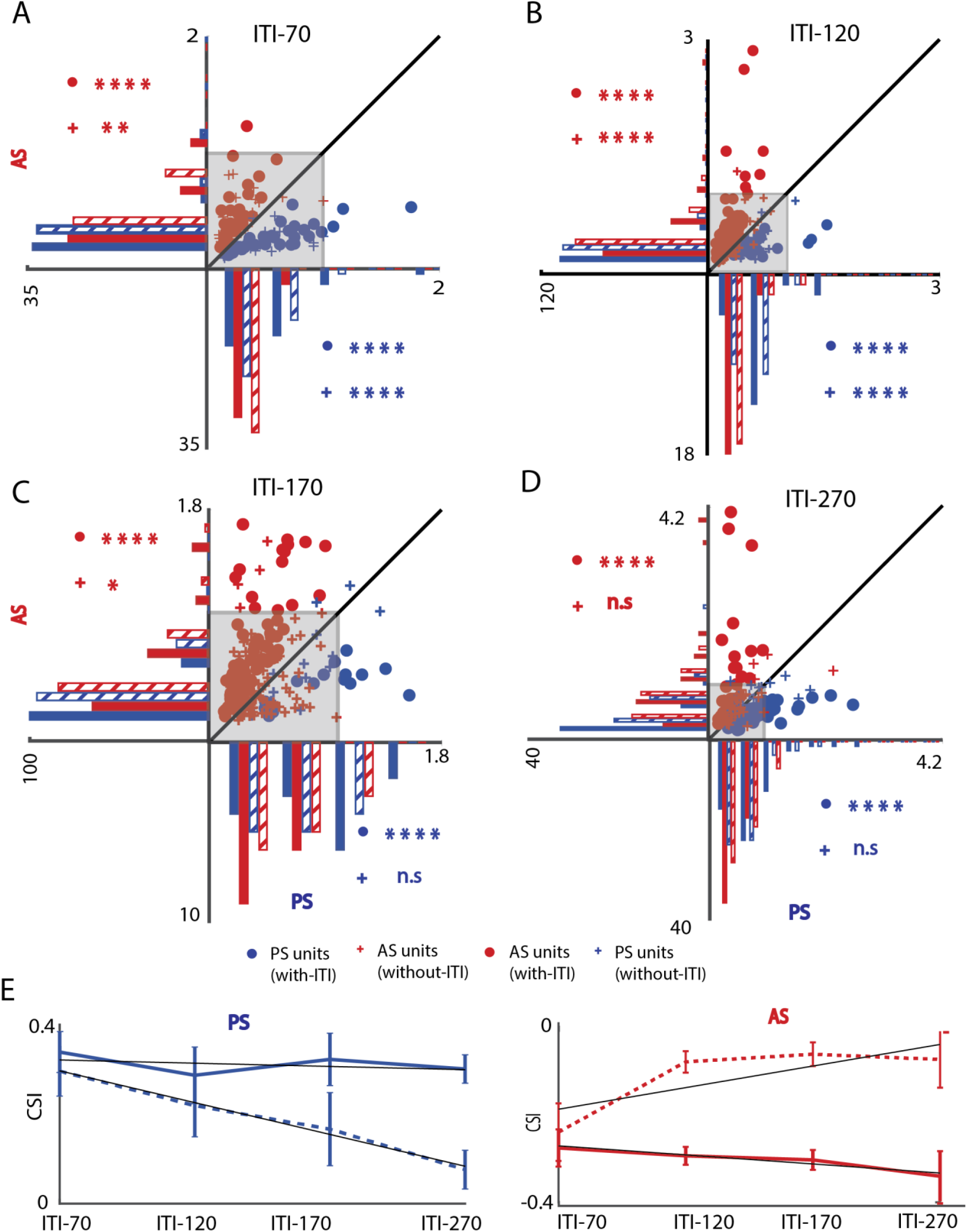
Enhanced responses emerge with longer ITIs. (**A-D**) Scatter plots showing the Normalized Cumulative Response Rates (*CRR*^*Norm*^) for Periodic-Selective (PS) and Aperiodic-Selective (AS) subpopulations for each unit, comparing periodic and aperiodic stimuli across all multiple-ITI stimulus sets, ITI-70, ITI-120, ITI-170, and ITI-270, respectively. Filled circles represent responses with ITI, while ’+’ symbols represent responses without ITI. The x-axis represents the normalized cumulative firing rates for PS units, while the y-axis represents the normalized cumulative firing rates for AS units. A 45° diagonal line is shown, where units below the line are categorized as PS, and units above the line as AS. Histograms along the axes display the distribution of *CRR*^*Norm*^ with filled bars for responses with-ITI and dashed bars for responses without-ITI. Significant differences in *CRR*^*Norm*^is marked with asterisks: •* = with-ITI; +* = without-ITI. Colors denote subpopulations (PS, blue; AS, red). (Two-tailed t-test:, *p<0.05, **p<0.01, ***p<0.001, ****p<0.0001, and n.s. = Not significant). **(E**) Common Selectivity Index (CSI) values for Periodic-Selective (left) and Aperiodic-Selective (right) units under both with-ITI (solid lines) and without-ITI (dashed lines) for all multiple-ITI stimulus conditions. Error bars indicate 95% confidence intervals (bootstrap, n = 1000).

To assess the influence of ITI on neuronal selectivity as shown in Fixed-ITI stimulus set, we directly compared the difference in *CRR*^*Norm*^ between periodic and aperiodic sequences obtained with and without ITI. For each neuron, we computed *CRR*^*Norm*^ (Periodic ,with-ITI) - *CRR*^*Norm*^ (Aperiodic ,with-ITI) and *CRR*^*Norm*^ (Periodic ,without-ITI) - *CRR*^*Norm*^ (Aperiodic ,without-ITI) , and compared these two quantities using a t-test within PS and AS populations. The PS units showed significant effects of ITI (ITI-120, p=0.049; ITI-170, p=0.0014; ITI-270, p=4.7*10^-6^) and AS units showed differences (ITI-120, p=7.8*10^-9^; ITI-170, p=5.23*10^-10^; ITI-270, p=0.0018) other than ITI-70PS, p=0.056 and ITI-70AS, p=0.507.

To summarize the above observations, common selectivity index (CSI, *Methods-Common Selectivity Index*) was calculated for both with-ITI and without-ITI conditions, separately for Periodic-selective (PS) and Aperiodic-selective (AS) units to quantify differences in coding of Periodic and Aperiodic stimuli. CSI values were consistently higher (and lower) for PS (and AS) subpopulations when the entire sequence, with-ITI (Figure 8E, blue and red solid lines), was considered compared to the without-ITI condition (Figure 8E, blue and red dashed plots). The result indicates enhanced selectivity for the preferred stimulus when silence durations were included. Although CSI remains similar for PS (positive values) and AS (negative values) units across different ITIs, removing the ITI period activity made CSI for both unit types go towards 0 with increasing ITI (Figure. 8E, dashed lines).

### Average rate responses of the population of neurons do not show selectivity except in offset at long ITIs

Mean responses of single units to Periodic and Aperiodic stimuli were calculated separately for each segment (300 ms, Period-3 or 400 ms, Period-4) across all trials, with 95% confidence intervals estimated using bootstrapping (*n* = 1000). The responses for Periodic stimuli were the same as Aperiodic stimuli in fixed-ITI stimulus set, with overlapping error bars indicating no clear separation in population mean firing rate throughout the stimulus (Figure. 9A). Across all ITI durations in multiple timescale-ITI stimulus set also, the averaged response profiles showed no significant differences in encoding patterns between Periodic and Aperiodic stimuli at the population level (Figure. 9B). This indicates that when considering all neurons collectively, no significant encoding differences emerge, regardless of stimulus type or ITI variability.

**Figure 9.**
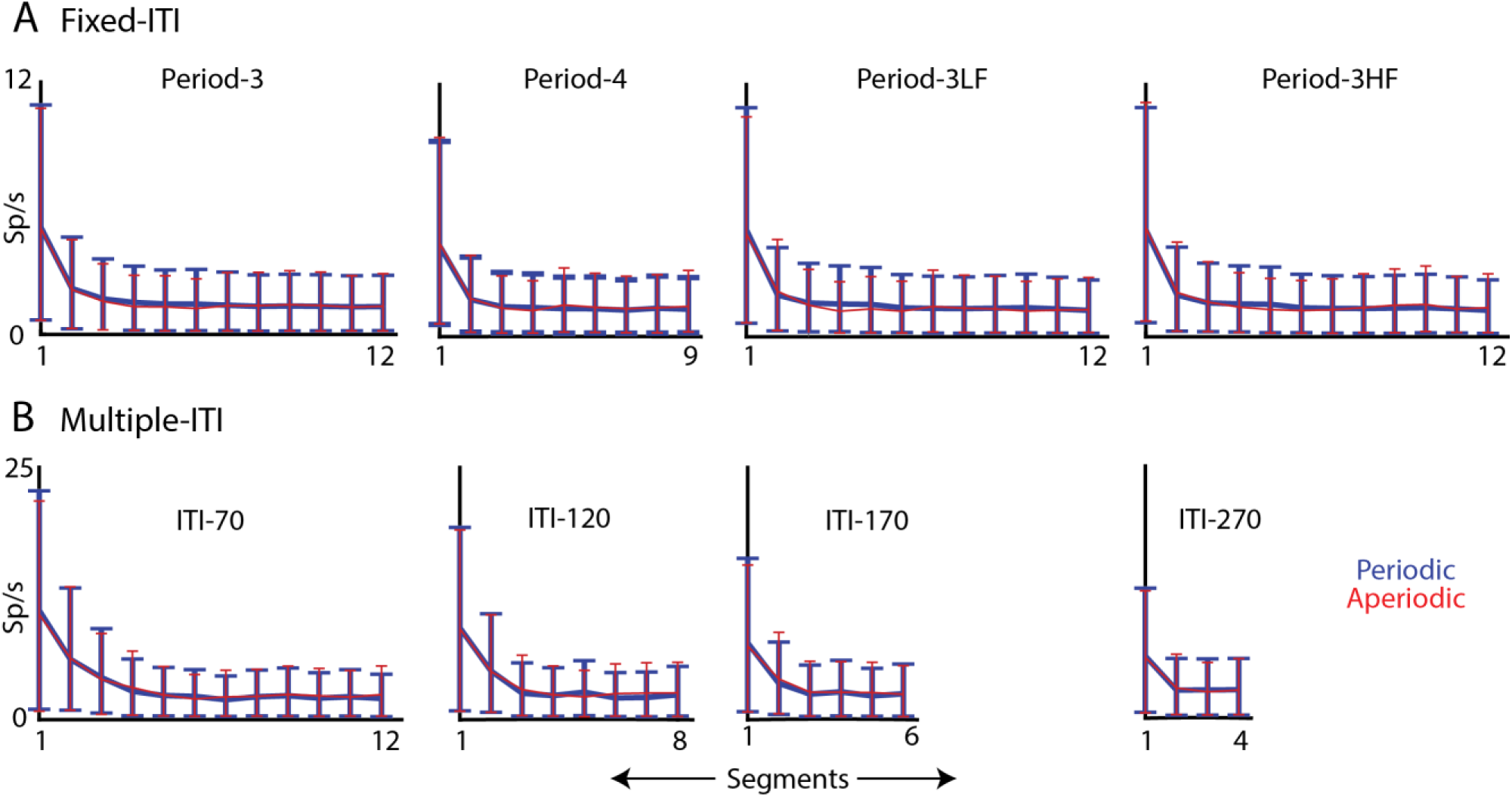
Lack of difference in population response to periodic and aperiodic stimuli. **(A)** Mean firing rates of all single units for periodic (blue) and aperiodic (red) stimuli in the fixed-ITI conditions, Period-3, Period-4, Period-3LF and Period-3HF respectively, segmented by stimulus cycle (300 ms for Period-3, 400 ms for Period-4). **(B)** Same as (A), but for multiple-timescale ITI conditions (ITI-70 to ITI-270). Error bars indicate 95% confidence intervals (bootstrap, n = 1000).

Offset responses were assessed for multiple -ITI stimulus set incorporating all trials (5 trials each for baseline and offset) across all 306 single units by finding the differences in rates Periodic and Aperiodic stimuli with same spectral content across 1ms bins, as 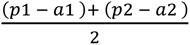, where *p*1 and *a*1 is the average response across all trials for PHF and AHF stimulus sets and *p*2 and *a*2 for the PLF and ALF (*Methods-Auditory Stimulus with multiple timescales)*. Significant differences were identified using paired t-tests, where 400 ms before stimulus onset served as the baseline and 400 ms after stimulus end as the offset response window. At ITI- 70 (p=0.31) , and ITI-120 (p=0.08) showed no offset effect, whereas ITI 170 (p=0.00019) and ITI 270 (p=0.009) showed significant offset based information to discriminate periodic and aperiodic stimuli. For fixed-ITI stimulus set, with 70 ms ITI, as above did not show any significant offset based differences.

### ITI neuronal activity builds up during the stimulus providing periodic-aperiodic information

The periodicity-aperiodicity related information in the period after offset of the stimulus suggests that the same could be building up during the stimulus sequences over each segment or period during the interval between segments or the stimulus off period of the previous segment (Figure 10A, left). Thus, we considered the average responses in the 50 ms interval preceding each segment of the stimuli (A1, P1, A2, A3, P3, …, Figure 10A). The difference between Periodic and Aperiodic conditions were obtained, as before τ𝑒𝑙*R* = (𝑝1 − 𝑎1 ) + (𝑝2 − 𝑎2 ) (*Methods-Auditory Stimulus with multiple timescales)*, for the above mentioned 50 ms interval for each segment. A linear fit (*polyfit* function in MATLAB) was made to the population average rate to account for changes along the stimulus for every segment (Figure 10A, right). The slope mean estimate and its 95% confidence interval were obtained by bootstrap (*n*=500) with randomly resampling with replacement from the population of neurons. Positive values of the slope would mean buildup of a predictive signal before the stimulus segment.

**Figure 10.**
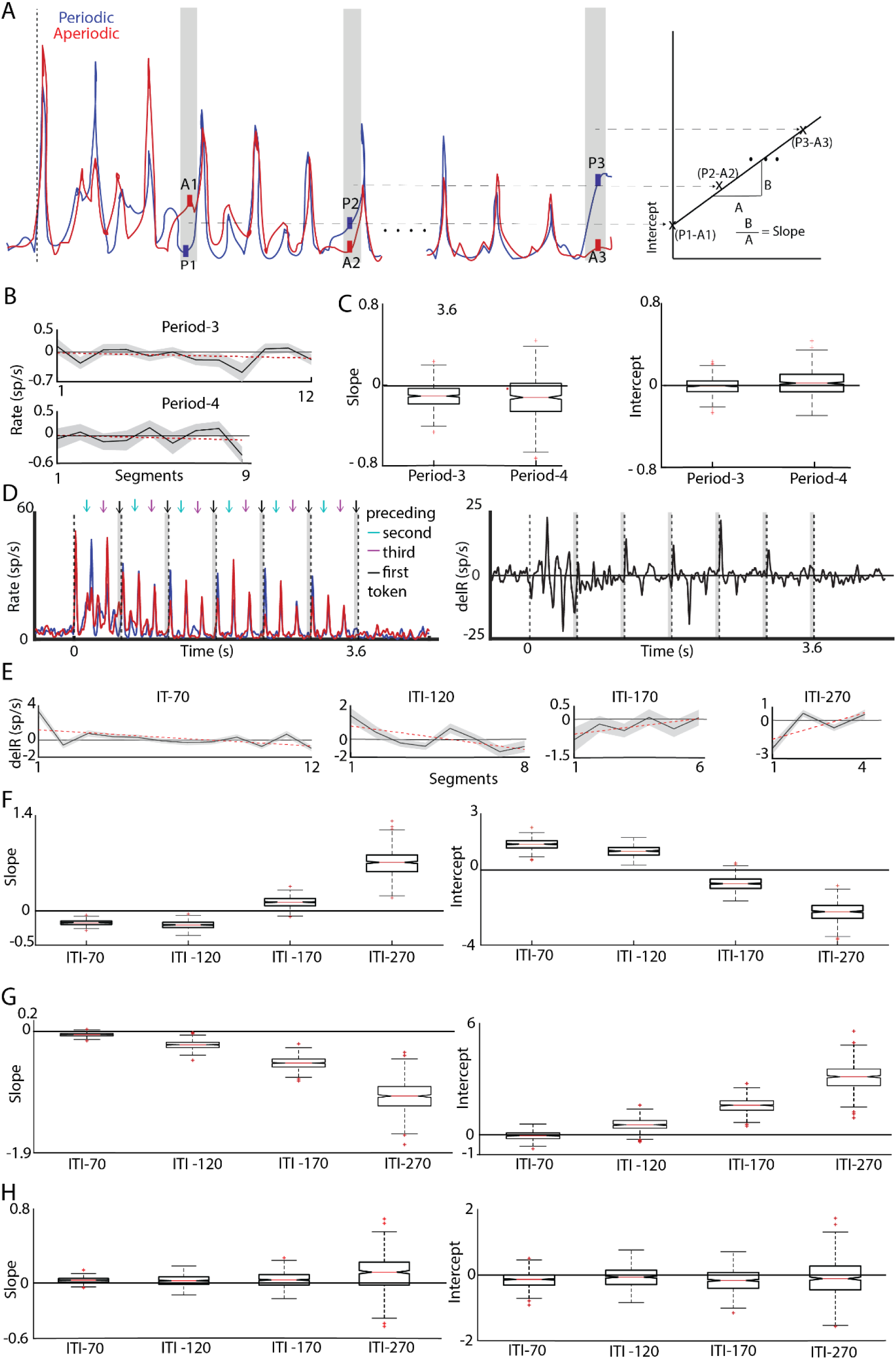
Buildup of prediction related information during the stimulus sequence. **A (Left)**: Schematic responses to Periodic (blue) and Aperiodic (red) stimuli aligned to individual segments (P1, A1, P2, A2, etc.). Gray shaded areas indicate 50 ms windows preceding each segment used for analysis. **(Right)** Schematic showing linear slope fitting over the pre-segment intervals based on difference scores with slope and intercept estimated via linear regression (MATLAB polyfit). **(B)** Mean difference in population response (Periodic – Aperiodic) in the 50 ms before each stimulus segment (300 ms in Period-3 and 400ms in Period-4 fixed-ITI stimulus set. Gray shaded regions represent the standard error of the mean (SEM), and red dashed lines indicate linear fits across segments. **(C)** Boxplots of slope and intercept values obtained from bootstrap resampling (n = 500) for Period-3 and Period-4 stimulus set. **D (Left)** Example response patterns of periodic (blue) and aperiodic (red) stimulus set averaged from 10 single units showing the 50 ms intervals immediately preceding the onset of each token in a segment (black arrows: first token; cyan and magenta arrows: second and third tokens) from the ITI-170 set. **(Right)** Response differences (delR) between Periodic and Aperiodic stimuli, averaged across the same 10 units as in the left panel, showing 50 ms intervals before each segment (gray shaded). **(E)** Mean difference in population response (delR) between periodic and aperiodic stimulus in the 50 ms before each stimulus segment across all multiple-ITI stimulus sets (ITI-70, ITI-120, ITI-170 and ITI-270 respectively). Gray shaded regions represent the standard error of the mean (SEM), and red dashed lines indicate linear fits across segments. **(F)** Boxplots of slope (left) and intercept values (right) for each ITI condition from bootstrapped population (E) fits preceding the first token. **(G)** Boxplots of slope and intercept values for each ITI condition preceding the second token. **(H)** Boxplots of slope and intercept values for each ITI condition preceding the third token. Each boxplot reflects 500 bootstrap iterations with 95% confidence intervals.

The difference in average responses to Periodic stimuli and its aperiodic counterparts in the 50ms intervals preceding each period (Figure 10B) do not show a consistent trend or prediction for either Periodic 3 or Periodic 4 in fixed-ITI stimulus set (Figure 10C). The slopes remain near zero (Period-3, −0.11*10^-4^; Period-4, −0.13*10^-4^), suggesting no significant trend in response magnitude over time. Similarly, the intercept values cluster near zero (Period-3 , −0.09* 10^-4^ ; Period-4, 0.25*10^-4^) indicating no consistent baseline response that differentiates between different Periodicities. Fixed-ITI stimuli with different spectral content showed slope values (Period-3LF, 0.04* 10^-3^; Period-3HF, 0.11*10^-3^) and the intercept values (Period-3LF, 0.18*10^-3^; Period-3HF, −0.1*10^-3^) does not indicate any predictive trends.

Next we considered responses to Periodic and Aperiodic stimuli with different ITIs (Figure 10D, left for ITI = 170 ms). From the differences in the PSTHs (Figure 10D, right), the (𝑝1 − 𝑎1 ) + (𝑝2 − 𝑎2 ) the successive differences in the mentioned duration before each segment (gray bar in Figure. 10D) were fitted with a straight line, as with fixed-ITI stimuli (Figure. 10B), across segment number (Figure. 10A, right, Figure. 10E, black lines with 95% confidence intervals in gray) for each ITI (Figure 10E, red dashed line).

Analysis of multiple-ITI stimulus sets across all periods revealed distinct trends in predicting Periodic sequences, indicating variability in neuronal response patterns (Figure. 10E). For ITI-70 and ITI-120 the slope values considering all periods remain negative, close to zero [ ITI-70 (−0.2 * 10-3) ; ITI-120 (−0.2 * 10-3), Figure. 10F, left], indicating lack of predictive information in the shorter ITI conditions. For ITI 170 and ITI 270 the slope values are noticeably positive [ ITI-170 (0.12 * 10-3); ITI-270 (0.7 * 10-3), Figure. 10F, left], particularly for ITI-270, where the trend is more pronounced. Bootstrapped box plots (500 iterations) with 95% confidence intervals (Figure 10 F) revealed a pronounced trend for ITI-270, indicating a temporal increase in neuronal activity preceding each period. This suggests the presence of a predictive mechanism, emphasizing the role of ITI duration in shaping anticipatory neuronal responses and also in discriminating periodic and aperiodic sequences.

To check whether the above buildup occurs specifically preceding every period (first token of a segment) we analyzed the 50 ms windows preceding second and third tokens also (Figure. 10D, left, cyan and magenta arrows) in the same way as above. Slopes and intercepts of population neuronal response differences were calculated as before (Figure. 10A, right). The 50 ms interval before the second token in each period revealed notable slopes, indicating temporal changes in neuronal activity, suggesting predictive buildup beyond just the first token. The analysis reveals a negative trend in the slopes of neuronal responses preceding the second token as the ITI increases (Figure. 10G). The slopes are close to zero in the case of ITI −70 and ITI-120 (ITI-70, −0.14*10^-4^; ITI-120, −0.18*10^-3^), indicating minimal changes in neuronal responses before the third token while the longer ITIs (ITI-170, −0.5*10^-3^; ITI-270, −1.0*10^-3^) show a decreasing trend in response difference. The same analysis of the 50 ms windows before the third token of each period showed no buildup of information as slopes (ITI-70, 0.03 * 10^-3^; ITI-120, 0.03* 10^-3^, ITI-170, 0.04* 10^-3^; ITI-270, 0.11*10^-3^) and intercepts (ITI-70, −0.14* 10^-3^; ITI-120, −0.09* 10^-3^; ITI-170, −0.17* 10^-3^; ITI-270, −0.1* 10^-3^) were not significantly different from 0 (Figure 10H).

## Discussion

Majority of the literature on neural coding of sound sequence has focused on deviation from regularity, either as stimulus specific adaptation (Ulanovsky et al., 2003; Ulanovsky et al., 2004), as deviant detection (Nelken, 2014), as repetition suppression (Todorovic & Lange, 2012; Mayrhauser et al., 2014; Auksztulewicz & Friston, 2016) or prediction error (Parras et al., 2017; Audette & Schneider, 2023) in multiple species including humans (Takasago et al., 2020; Hu et al., 2024). The auditory system’s sensitivity to the above violations of regularity is important as it allows understanding of deviation from important regular sequences signaling unexpected events which can be important for the animal. Regular or predictable sequences, with the same auditory content repeating successively, are important in multiple contexts, like social communication in mice, experience dependent plasticity, auditory stream segregation, grouping and various aspects of auditory scene analysis (Perrodin et al., 2023; Agarwalla et al., 2020; Agarwalla et al., 2023; Agarwalla et al., 2025; Andreou et al., 2011; Carl & Gutschalk, 2012). Thus, coding of regular sound sequences in the auditory system deserves more attention than it has currently received.

The general observation is that, due to adaptation or repetition based suppression, responses to regular sequences weaken over time compared to irregular sequences. In a previous study (Mehra et al., 2022), a substantial proportion of layer 2/3 neurons in the ACx displayed a preference for regularity, whereas another subset showed a preference for irregularity when a deviant sound was introduced at either fixed or random intervals within an oddball paradigm. Contrary to the general notion of observing suppression for repeating stimuli, neurons could get enhanced in their responses to repeating stimuli as well. In the current study we expand on the above finding and have introduced a range of sound stream periodicities, frequency content and inter-token intervals to investigate how ACx neurons encode periodic and aperiodic stimuli.

In, both single unit and two-photon Ca^2+^ imaging data we identified two subpopulations of neurons that exhibit selectivity for Periodic or Aperiodic stimuli. This selective responsiveness was consistently observed across Periodicities and different inter token intervals (ITI) suggesting a robust, sequence type specific tuning in such neurons. The convergence of results from single units in anesthetized mice (Table 1 and Table 5) and single neuron resolution Ca^2+^ imaging in awake mice (Table 3) supports the functional significance of these subpopulations.

Our findings collectively indicate a lack of generalization in neuronal selectivity across varying Periodicities, spectral properties, and inter-token intervals. Negligible number of single neurons exhibited consistent responses spanning all stimulus categories based on periodicity or frequency content (Table 2 and Table 4), whether they were PS or AS. Further, primarily different subsets of single neurons were PS or AS across the different ITIs tested (Table 6).

With the Ca^+2^ imaging data we also observed a micro-organizational feature of such selectivity. We considered noise correlations between pairs of neurons and probability of observing EP at certain distance from EP and similarly EA from EA neurons. We observed that there is a heterogeneous connectivity network locally, with a large majority being NS often connected with themselves and also other types, or non-responsive, with 100 μm diameter clusters of multiple EA or EP neurons. Thus, the subpopulations of neurons observed here could form functional groups and may be a basic feature in ACx micro-organization. Further role of the observed noise correlations can be studied in discriminating periodic and aperiodic stimuli by the populations of neurons (Averbeck et al., 2006; Cohen & Maunsell, 2009). In these experiments, mice were passively listening to the stimuli and were naïve in terms of the types of stimuli. It remains to be seen how the above organization, EP and EA neuron proportions, discriminability of the stimuli by populations of neurons and generalization ability get altered if mice were trained and engaged in an albeit difficult task to differentiate such stimuli (Periodic and Aperiodic) in a general manner – across periodicity, frequency content or ITI.

Interestingly, we observed that some neurons displayed facilitation instead of adaptation for a sound sequence with tokens and periods repeated (Figure. 3 and Figure. 8A-D, units depicted outside gray boxes). Moreover, the profiles of normalized PS or AS population rate responses for each segment across the sound sequence (Figure. 2C-F and Figure. 7C-F) could also show little to no decrease of responses for Periodic stimuli or Aperiodic stimuli further emphasizing the robust nature of selectivity. In general, it was observed that with increasing ITI such enhancement increased. The significance of the ITIs used in our study is that they are within the main range of inter syllable intervals observed in mouse vocalization sequences which have been shown to be possibly coded as a whole or single object and not as individual tokens (Agarwalla et al., 2020; Agarwalla et al., 2023).

Most importantly we observed a profound role of neural activity during the interval between tokens. In the normalized rate response profiles as above, removal of the activity in the silence period could alter the temporal evolution of the response markedly. In most cases a stronger suppression is observed when only responses to tokens are included, leaving out the activity during the intervening silences (Figure. 2C-F and Figure. 7C-F). Such suppression was almost always true in case of responses of PS to Periodic stimuli and that of AS to Aperiodic stimuli. Effect of the ITI neural activity was the same as above, for PS when responding to Aperiodic stimuli and AS when responding to periodic stimuli except at the long ITIs (170 ms and 270 ms). The effect of the ITI neural activity was the most at the above ITIs, with maximum gain due to inclusion of the gap period activity in separating Periodic and Aperiodic stimuli achieved at 270 ms ITI (Figure. 8E). The above result suggests the sequential integration of the tokens into a sequence in the neural response and may underlie possible creation of a sound object.

Sound offset responses play a crucial role in auditory perception, particularly in processing continuous sounds such as speech (Liberman et al., 1954; Ghitza, 2011). They are integral to encoding sound duration (Malone et al., 2015; Li et al., 2021; Sołyga & Barkat, 2021) and detecting silent gaps between sounds (Anderson & Linden, 2016; Weible et al., 2020). Auditory cortex exhibits offset responses to repetitive click stimuli, modulated by both inter-click interval (ICI) length and the total number of clicks (Song et al., 2024) and help in temporal integration. Short gaps (2ms – 20 ms) in auditory stimuli also elicit distinct offset responses in the auditory cortex of unanesthetized rats (Awwad, Bshara et al. 2023). Offset responses were also shown to be involved in tying together 2 sound syllables (Lee & Rothschild, 2023) and carry information about preceding sounds in the auditory cortex (Lamothe et al., 2025). Similarly, here we find that although population mean rate responses do not show differences between periodic and aperiodic stimuli, but stimulus off period activity was different from baseline in different ways for Periodic and Aperiodic stimuli providing information about the stimulus type.

Our findings link cortical selectivity for periodic/aperiodic stimuli to intrinsic brain dynamics, where external periodic inputs may engage rhythmic oscillations and aperiodic inputs to broadband activity (Yu et al., 2024). This correspondence offers a promising avenue for acoustic decoding BCIs, as integrating these specialized streams via representation learning (Yu et al., 2025) can enhance decoding accuracy by learning distinct neural manifolds for periodic (e.g., temporal) and aperiodic (e.g., spectral) acoustic features.

Different preferences for periodic versus aperiodic sequences in auditory cortex likely arise from several interacting mechanisms: short-term synaptic plasticity that drives adaptation and context sensitivity (Reyes A. D. 2011; Li, J. Y., & Glickfeld, L. L , 2023), predictive-coding signals that build expectations and flag deviations (Barascud et al., 2016) , periodicity/pitch-tuned neurons (Bendor, & Wang 2010), and subtype-specific inhibition from PV (fast) and SOM (slower) interneurons (Natan et al., 2015; Natan et al., 2017). The differential selectivity we observe for periodic versus aperiodic sound sequences is likely rooted in the sophisticated computational architecture of the canonical thalamo-cortical microcircuit (Antunes, & Malmierca 2021). Specifically, the precise interplay between excitatory inputs, distinct inhibitory interneuron populations (PV, SOM, VIP) ((Studer et al., 2021; Blackwell & Geffen, 2017), and interlaminar processing dynamically shapes network responses, enabling the segregation of temporal regularity from unpredictable transients.

Based on the above observations we hypothesized that following each period in Periodic stimuli or an equivalent segment in Aperiodic stimuli, the intervening period till the next segment carries differential information for the Periodic and Aperiodic stimuli. Thus, as the stimulus progresses, neural activity in such intervening periods may act as an anticipatory signal for the next segment and can play a role in prediction error signals. Since these are in anesthetized mice, top down signals are not engaged and still we may have prediction error signals based on feed forward stimulus activity and local circuit properties. Presence of omission responses in anesthetized rodents (Lao-Rodriguez et al., 2023) can be explained based on such an anticipatory signal. We show that such a signal differentiating the two types of stimuli builds up during the stimulus, and is stronger as the ITI is increased. The mechanisms underlying such signals could be differential adaptation of inhibitory neuron types and release from inhibition, which can be deciphered with future optogenetics experiments.

## Limitations of the Study

Our study offers valuable insights into the lack of generalization of selectivity to periodicities, spectral contents, and inter-token intervals in a sound sequence. It also shows prediction to periods with increasing inter-token intervals. However, the order of pure tone arrangements in our sequences (F1–F2–F2 or F2–F1–F1) makes it unclear how spectral arrangement may influence neuronal responses. Naturalistic sounds or species-specific vocalizations may engage different neural circuits, altering the selectivity patterns. Furthermore, the calcium imaging technique employed here is inherently limited in temporal resolution, potentially missing fast or transient neural dynamics critical for encoding temporal structure. Another concern is the absence of behaviorally relevant context or task engagement, where attention-or expectation-driven responses are not considered. Future studies incorporating behavior, natural sounds, and improved temporal resolution are necessary to address these limitations.

## RESOURCE AVAILABILITY

### Lead contact

Further information and requests for resources should be directed to and will be fulfilled by the lead contact, Dr. Sharba Bandyopadhay.

### Materials availability

This study did not generate new unique reagents.

### Data and code availability

All custom analysis code, are publicly available in Mendeley Data. https://data.mendeley.com/drafts/ddt89mbtxt. micheal, ann soniya (2025), “Periodic and Aperiodic sound sequences-A1-codes”, Mendeley Data, V1, doi: 10.17632/ddt89mbtxt.1

## Acknowledgments

ASM thanks MoE for Institute Fellowship, SB thanks IIT Kharagpur and SRIC Cell, IIT KGP for start-up and Challenge Grant funds. This work was supported by India-Czech Bilateral Scientific and Technological Cooperation DST India DST/INT/CZ/P-04/2020, SERB-CRG/2021/005653 and DST/CSRI/2021/340 awarded to SB.

## Author Contributions

ASM and SB conceived the project, ASM performed all experiments, ASM and SB analyzed data. SB supervised the work. ASM, SB wrote the manuscript.

## Methods

### EXPERIMENTAL MODEL AND STUDY PARTICIPANT DETAILS

#### Animals

Experiments were conducted in mice (Mus musculus), encompassing both sexes and aged between 8 to 12 weeks, with the approval of Institutional Animal Ethics Committee (IAEC) of Indian Institute of Technology Kharagpur. Across all experiments 14 mice served as experimental subjects. Animals were maintained under a 12/12 h light/dark cycle at a temperature of 25°C and had access to food and water *ad libitum*. In-vivo extracellular electrophysiology experiments were performed on NIN Hyderabad stock animals and two-photon calcium imaging on C57BL/6J-Tg (Thy1-GCaMP6f, strain #024339 , The Jackson Laboratory).

#### Animal Preparation for acute In vivo extracellular electrophysiology

Prior to the initiation of surgery, mice were placed under profound anesthesia using 5% isoflurane, with the anesthesia levels subsequently maintained at 1.5–2.5% and body temperature at 37°C for the duration of both the surgical procedure and the ensuing experiment. An incision was done along the midline to expose the left temporal area of the scalp. This area was then cleared of tissue, and sterilized by the application of 3% hydrogen peroxide (H_2_O_2_), followed by 70% alcohol. A titanium head plate was fixed on the skull using dental cement to head-fix the animal during the experiment. A circular craniotomy of ∼2mm was made with a microdrill, over the presumed left auditory cortical area, which was demarcated by the temporal ridge, the lambdoid suture, and both ventral and rostral squamosal sutures (Mehra et al., 2022a). These animals were subjected to acute in vivo electrophysiology.

#### Animal Preparation for chronic 2-photon Ca^2+^ imaging

All procedures followed up to the headplate fix remain the same for the chronic awake 2-photon Ca^2+^ imaging. The head plate is fixed using a meta bond and a circular craniotomy of ∼3mm is made using microdrill over the left auditory cortical area. The exposed cranial window was covered with a 3 mm coverslip. The animal received antibiotics for five days following the surgery; after recovering, the animal was trained on the imaging platform for five days to prepare for awake recordings.

### METHOD DETAILS

#### Auditory Stimulus

##### Auditory Stimulus with fixed time scale - different Periodicities (fixed-ITI)

The fixed time scale stimuli consisted of a sequence of 36 tokens of pure-tones, each 30 ms long with 5 ms rise and fall times followed by 70 ms silence (Figure. 1A). The tokens were one of two distinct frequencies (*f_1_* and *f_2_*, Figure. 1, blue and red bars) chosen based on the best frequency (BF) distribution of single units recorded simultaneously (see below in “*In-vivo extracellular recordings*”). *f_1_* and *f_2_* were separated by 1.3 octaves and generally *f_1_* was near or at the majority of BFs of units encountered in a particular recording location. The stimuli were either Periodic (Figure. 1A, left) or Aperiodic (Figure. 1A, right) and further categorized based on their Periodicities. Either 3 or 4 successive tokens were repeated to create Periodic stimuli and thus had Periodicities of 3 or 4 tokens referred to as Period-3 or Period-4 respectively (Figure. 1A, top two sets, and bottom set, left). Period-4 stimuli always had 2 tokens each of *f_1_* and *f_2_*, while Period-3 stimuli were categorized based on the number of *f_1_* and *f_2_* tokens in each period. Period-3 stimuli with more of the higher frequency tokens present are referred as Period-3HF (one *f_1_* and two *f_2_* tokens) and the others as Period-3LF (two *f_1_* and one *f_2_* tokens). For the analyses we considered 4 groups: Period-3 (ALL, consisting of both the next two groups), Period-3HF (f1-2f2, Figure. 1A, top set, left), Period-3LF (2f1-f2, Figure. 1A, middle set, left), and Period-4. Each group of Period-3 stimuli included three Periodic stimuli (Figure. 1A, P3HF and P3LF, left, top 2 groups) and two Aperiodic stimuli (Figure. 1A, right, top 2 groups A3HF and A3LF). A3HF and A3LF were created such that the number of tokens in each period of P3HF or P3LF corresponded. The Period-4 category, contained 4 Periodic stimuli (Figure. 1A, bottom group, left) and 2 Aperiodic stimuli (Figure. 1A, bottom group, right). Thus, in total, the fixed ITI set comprised of 10 Periodic and 6 Aperiodic sequences. Each sequence includes 36 pure tone tokens and 36 silence periods. Each sequence is repeated five times in a pseudorandom order, with a 2-second inter-stimulus interval. Single-unit spiking responses are recorded with a 500 ms baseline. These stimuli were presented at 60-70 dB SPL.

##### Auditory Stimulus with multiple timescales

Organized into four sets, the stimuli were differentiated by the Inter token intervals (ITI), varying systematically across the sets at 70 ms, 120 ms, 170 ms and 270 ms respectively. As in the above fixed-ITI stimulus set, each pure tone token in the multiple timescales or multiple-ITI stimulus set is of 30 ms duration, with 5 ms rise and fall times and two different frequencies (*f_1_* and *f_2_*, 1.3 octaves apart). The total duration of each stimulus, regardless of the ITI, was kept constant at 3.6 seconds and played at 60-70 dB SPL. There were 4 stimuli corresponding to each ITI, all with a period of 3 tokens, featuring both Periodic and Aperiodic sequences (Figure. 2B). Each stimulus includes two different frequencies, as in the fixed-ITI stimulus set. The 4 stimuli in each ITI set above, consisted of 2 Periodic and 2 Aperiodic stimuli having 1 *f_1_*-token and 2 *f_2_*-tokens (PHF), and 2 *f_1_*-tokens and 1 *f_2_*-token (PLF) and one realization of their Aperiodic counterparts (AHF and ALF). Thus, the multiple-ITI stimulus set consisted of a total of 16 stimuli, 8 Periodic and 8 Aperiodic. The stimuli were presented in a pseudorandom order from the set of 16 and repeated 5 times, with an inter-stimulus interval of 2 s, and single-unit spiking with 500 ms baseline was recorded.

##### Stimulus Delivery (Electrophysiology and 2-photon Ca^2+^ imaging)

Sound stimuli were digitally generated using custom software written in MATLAB (MathWorks) and were produced by NI Data Acquisition modules. These signals were then delivered via a TDT RX6, attenuated through a TDT PA5 attenuator, and presented through electrostatic speakers (ED1). During two-photon Ca^2+^ imaging experiments, the same stimuli were delivered using a TDT RZ6 high-frequency auditory signal processor. The speakers were positioned approximately 10 cm from the contralateral ear. Measurements of the ES1 speaker’s frequency response, taken with a 4939 microphone (Bruel & Kaer), revealed a typically flat (+/-7 dB) calibration curve extending from 4 to 60 kHz.

#### In Vivo Extracellular recordings

Extracellular recordings were conducted in vivo utilizing a 4 × 4 Tungsten multielectrode array with a 125-μm interelectrode spacing and an impedance of 3–5 MΩ (MicroProbes). The electrodes were precisely positioned to probe the responses in layer 2/3, at a depth ranging from 200-300 𝜇𝑚 from the cortical surface, with the assistance of a micromanipulator (MP-285, Sutter Instrument Company). The electrodes were delicately inserted into the targeted layer 2/3 and allowed to settle for a period of 15 minutes prior to initiating the recording process. The acquired signals were initially passed through a unity gain (1×) head stage, subsequently amplified by a preamp (Plexon, HST16o25) with a gain factor of 1000. The system simultaneously recorded wideband signals, encompassing local field potentials (ranging from 0.7 Hz to 6 kHz), and spike signals (within the frequency range of 150 Hz to 8 kHz). This data acquisition was facilitated through a National

Instruments Data Acquisition Card (NI-PCI-6259), operating at a sampling rate of 20 kHz. Subsequent analysis of the obtained signals, either online or offline, was conducted using custom codes developed in MATLAB (MathWorks Inc.). For each stimulus delivery after 500 ms, the data acquisition lasted up to 5s.

##### In vivo 2-photon Ca^2+^ imaging

Two-photon calcium imaging was performed on a Bruker corporation laser scanning microscope using Prairie View software (Prairie View Technologies). The excitation beam of wavelength 860 nm generated with a Spectra Physics Mai Tai Deep See Ti-Sapphire mode-locked femto-second laser and was delivered through a 20X 0.8 NA water immersion objective (2 mm WD, Olympus). Each frame in the ROI ∼228 𝜇𝑚 × 137𝜇𝑚, with pixel size 1.17 𝜇𝑚 was obtained at a frame period of ∼196 ms. Frames were acquired at a depth of 200-300 𝜇𝑚 with a frame period of ∼5fps with the stimulus delivered at the 9th frame for a 40-frame cycle of acquisition.

### QUANTIFICATION AND STATISTICAL ANALYSIS

#### Data Processing -Electrophysiology

##### Spike sorting

Spike sorting was performed using custom MATLAB scripts (MathWorks Inc.). Power supply fluctuations were removed by a 50 Hz notch filter. Spikes from 16 channels that were above a threshold of four standard deviations were extracted from the raw data. These spikes were then placed into a space defined by the top three principal components. After identifying the single-unit responses by calculating the isolation distances between cluster centers using K-means clustering, we checked the quality of each spike waveform by careful visual inspection. Using principal component analysis and K-means clustering, we carefully extracted the spike timings for all the channels.

##### Significant units’ selection

After spike sorting, units were deemed significant based on an unpaired t-test. For every stimulus set (fixed-ITI and multiple-ITI), a unit was considered responsive if there was a significant difference (with Bonferroni correction) between the firing rates during the baseline period (450–500 ms before stimulus onset) and the response period (500–550 ms after stimulus onset). A unit was classified as significant only if the set of 8 stimuli beginning with *f1*, as well as the set of 8 stimuli beginning with *f2*, each showed a response rate significantly different from baseline.

##### Normalized Cumulative response rates

To analyze the cumulative rate responses, we considered each stimulus as consisting of 𝑛 periods each period is composed of tokens and ITI. For the 𝑘^𝑡ℎ^ segment, the cumulative response rate *R*_*k*_ is defined as the average firing rate from the stimulus onset to the end of the 𝑘^𝑡ℎ^ period 𝑘𝑇, where 𝑇 is the duration of one period.

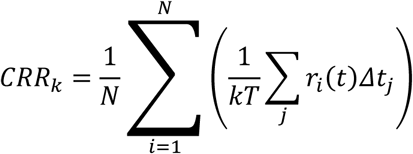

where, 𝑟_𝑖_(𝑡) represents the response rate for trial and 𝑁 is the total number of trials, 𝛥𝑡_𝑗_ is the time interval corresponding to each response measurement. These cumulative rates were then normalized by the first period rate( *R*_1_).

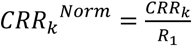 for *k* =1,2,…*n* (where 𝑛 is the total number of repetitions of the period)

For any given segment, the cumulative response rate is defined as the average firing rate from the stimulus onset to the end of that period. These rates are calculated by averaging the response rates across all trials. The overall normalized cumulative response rate of a stimuli for the entire sequence is denoted by *CRR*^*Norm*^.

##### Classification of single units

Single units that responded to Periodic and Aperiodic stimuli were classified into two categories: Periodic selective units (PS) and Aperiodic selective units (AS). This classification was applied to both fixed-ITI and multiple-ITI stimulus set. An unpaired t-test was conducted on the normalized mean cumulative rates (*CRR*^*Norm*^) of the entire stimulus duration for the Periodic stimulus sets [P3, P4, P3HF, P3LF) and the Aperiodic stimulus sets (A3, A4, A3HF, A3LF) for each single unit to determine their responsiveness to Periodic and Aperiodic stimuli.

If the *CRR*^*Norm*^ showed a significant effect favoring the Periodic set (right-tailed t-test) or the Aperiodic set (left-tailed t-test), the corresponding single units were classified as Periodic selective units (PS) or Aperiodic selective units (AS). A similar approach was applied to the multiple-ITI stimulus set. One-tailed unpaired t-test (p < 0.05), was conducted on the normalized mean cumulative response rates (*CRR*^*Norm*^) for the entire stimulus duration of all Periodic stimulus trials against all Aperiodic stimulus trials. The sound-responsive extracellular units were classified as Periodic selective (PS) or Aperiodic selective (AS) for each of the four fixed time scale stimulus categories based on the selectivity criteria.

##### With-ITI and Without-ITI

To understand the effect of silence in sound sequences, we analyzed response rates both with and without the silence duration. For stimuli composed of 30 ms tones followed by variable silence intervals (70 ms for fixed-ITI) we considered the response rates in two ways. First, by including the entire silence duration (With-ITI), we examined the full sound sequence. Second, excluding the last 50 ms of activity during 100 ms segment for each token (30 ms stimulus duration + 70 ms ITI). The 20 ms period following each tone was considered part of the response to the tone due to ACx latency.

In the case of the multiple-ITI stimulus set, a 20 ms duration following each tone (30 ms) was considered, as described above, while the remainder of the silence period in each token was excluded. For the multiple-ITI (Without-ITI), the silence periods of 50 ms, 100 ms, 150 ms, and 250 ms, corresponding to ITI-70, ITI-120, ITI-170, and ITI-270, respectively, were entirely excluded.

The influence of activity during the ITI was analysed using a parameter *d* defined as:

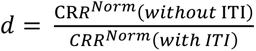. This parameter was calculated for a specific stimulus set (Periodic and Aperiodic) as an average of PS and AS subpopulations.

##### Common Selectivity Index

Common Selectivity Index (CSI), calculated as 𝐶𝑆𝐼 = (𝑝 − 𝑎)/(𝑝 + 𝑎) where *p* and *a* indicate the *CRR*^*Norm*^ for Periodic and Aperiodic stimuli respectively, within each ITI stimulus set. CSI was computed for all trials, averaged, and bootstrapped 1000 times to estimate 95% confidence intervals.

##### Data Processing - 2-photon 𝑪𝒂^(𝟐+)^ imaging

###### Significant cells selection

For each neuron, df/f was computed relative to baseline, defined as the mean fluorescence over frames 4–8 prior to stimulus onset. To determine whether a neuron was responsive, baseline activity was compared with stimulus-evoked activity across cumulative segments of the response window. Specifically, starting at stimulus onset (frame 9) and extending to frame 38, from the 40-frame cycle of acquisition, we constructed 10 cumulative segments, each formed by progressively aggregating 3 additional frames (∼600 ms). Thus, segment 1 included frames 9–11, segment 2 included frames 9–14, segment 3 included frames 9–17, and so forth, up to segment 10 (frames 9–38). For each of the 16 fixed-ITI stimuli, baseline df/f values (frames 4–8) were compared with each cumulative segment using an unpaired t-test. A Bonferroni correction factor of 16 × 10 = 160 was applied to control for these multiple comparisons within each neuron. A neuron was deemed responsive if it showed a significant difference in any segment (p < 0.05 after correction). Using this criterion, we identified 1,821 responsive neurons out of 2,319 total neurons, across 27 ROIs.

###### Classification of neurons

Sound-responsive neurons (Methods-significant cell selection) were identified through a comparative analysis of their mean baseline frame df/f with responses for fixed-ITI stimulus set. The mean df/f (9 to 33 frames, post stimulus 1 s) across all trials of a neurons to a particular Periodic stimulus set and its Aperiodic counterpart is compared to its baseline using unpaired t-test to get the significant neurons for Periodic and Aperiodic stimulus set. Then we examine if the neurons significant to Periodic stimuli also showed significance to its Aperiodic stimulus counterpart, and vice versa. Neurons exhibiting responsiveness to both Periodic and Aperiodic stimuli within a given Periodicity set were excluded from further analysis. This process was conducted independently for all fixed-ITI stimulus groups (Period-3, Period-4, Period-3HF, Period-3LF), resulting in two distinct populations of selective neurons. Exclusive Periodic neurons (EP) that are exclusively selective to Periodic stimulus set and Exclusive Aperiodic neurons (EA) that are exclusively selective to Aperiodic stimulus set.

###### Noise correlation

Pairwise noise correlations were computed across various neuronal categories (EP and EA) based on their stimulus selectivity. To analyze noise correlations, we first calculated the mean fluorescence change (df/f) across all frames from stimulus onset to 1 s after stimulus end for each trial denoted ^by 𝑟^_(𝑖,𝑠)_

Let 𝑟_(𝑖,𝑛)_represent the fluorescence change for the 𝑖^(𝑡ℎ)^ neuron in the 𝑛^𝑡ℎ^ trial of a particular stimulus,𝑠. We then subtracted the mean response of that stimulus across all trials from each individual trial response. The normalized response for the 𝑛^𝑡ℎ^ trial is 𝑟𝑠_(𝑖,𝑛)_) = 𝑟_(𝑖,𝑛)_ − 𝑟_(𝑖,𝑠)_, where 𝑟𝑠_(𝑖,𝑛)_is the noise vector for a particular stimulus. Similarly, noise vectors are calculated for all stimuli for a neuron and the Pearson correlation coefficient is calculated between the 𝑖^(𝑡ℎ)^ and 𝑗^(𝑡ℎ)^neuron in each category.

###### Spatial arrangement

The distance between type X (EP/EA) type neurons and the nearest Y type (EP/EA/NS(Non-selective)) was calculated. These distances were averaged across all ROIs after bootstrapping 500 times, to determine the mean raw distance between type X and type Y neurons. The probability of finding an X category neuron within distance of Y category was calculated. The mean distances between two categories of neuron types were considered significantly different if their 95% confidence intervals did not overlap.

